# Gra-CRC-miRTar: The pre-trained nucleotide-to-graph neural networks to identify potential miRNA targets in colorectal cancer

**DOI:** 10.1101/2024.04.15.589599

**Authors:** Rui Yin, Hongru Zhao, Lu Li, Qiang Yang, Min Zeng, Carl Yang, Jiang Bian, Mingyi Xie

## Abstract

Colorectal cancer (CRC) is the third most diagnosed cancer and the second deadliest cancer worldwide representing a major public health problem. In recent years, increasing evidence has shown that microRNA (miRNA) can control the expression of targeted human messenger RNA (mRNA) by reducing their abundance or translation, acting as oncogenes or tumor suppressors in various cancers, including CRC. Due to the significant up-regulation of oncogenic miRNAs in CRC, elucidating the underlying mechanism and identifying dysregulated miRNA targets may provide a basis for improving current therapeutic interventions. In this paper, we proposed Gra-CRC-miRTar, a pre-trained nucleotide-to-graph neural network framework, for identifying potential miRNA targets in CRC. Different from previous studies, we constructed two pre-trained models to encode RNA sequences and transformed them into de Bruijn graphs. We employed different graph neural networks to learn the latent representations. The embeddings generated from de Bruijn graphs were then fed into a Multilayer Perceptron (MLP) to perform the prediction tasks. Our extensive experiments show that Gra-CRC-miRTar achieves better performance than other deep learning algorithms and existing predictors. In addition, our analyses also successfully revealed 172 out of 201 functional interactions through experimentally validated miRNA-mRNA pairs in CRC. Collectively, our effort provides an accurate and efficient framework to identify potential miRNA targets in CRC, which can also be used to reveal miRNA target interactions in other malignancies, facilitating the development of novel therapeutics.

## Introduction

Colorectal cancer (CRC), or bowel cancer, occurs in the colon or the rectum. CRC is the third most common malignancy and the second leading cause of cancer death worldwide^1^. In 2020, there were approximately 153,000 new cases of CRC were diagnosed, and 52,500 deaths from CRC occurred in the United States^2^. There has been a notable increase in incident cases of colorectal cancer, with over 16 out of 21 global regions experiencing a doubling or more in cases in the past three decades^3^. By the year 2030, the global burden of CRC is expected to increase by 60%, involving over 2.2 million new cases and 1.1 million annual deaths^4^. This escalation is attributed to multiple factors, including the economic advancement of transitioning and low-to-medium Human Development Index nations, as well as shifts in societal norms within developed countries^5^. The rise in CRC incidence appears to be proportionate to economic development levels. This trend is thought to be driven by alterations in the environment and lifestyles, such as increasingly sedentary lifestyles, rising obesity rates, greater consumption of processed foods, alcohol, and meat, as well as overall increased life expectancy^6,7^. Although therapeutic approaches to treat CRC have improved in the past decade, both the incidence and mortality rates of CRC in adult patients below the age of 50 have increased by 22% and 13%, respectively^8^.

Specifically, about a quarter of CRC patients were diagnosed at an advanced stage, where the cancer has metastasized, most often to liver^9^. Metastatic CRC is associated with a particularly poor prognosis. Therefore, it is critical to probe the molecular determinants in CRC initiation, progression and metastasis to allow early detection and therapeutic intervention. Numerous studies have consistently detected aberrant expression of microRNAs (miRNAs) in CRC tissue and cells, suggesting that miRNAs play important roles in CRC^10^. The miRNAs, approximately, ∼22 <u>n</u>ucleotide (nt) in length, are ubiquitous gene regulators that modulate a broad range of essential cellular processes at the post-transcriptional level. Most miRNAs function in the cytoplasm, where they associate with <u>A</u>r<u>go</u>naute (AGO) proteins. AGO-miRNA complexes regulate target <u>m</u>essenger RNAs (mRNAs) through imperfect base-pairing with sequences in the 3′ <u>u</u>ntranslated region (UTR) to repress translation and cause mRNA deadenylation and decay^11,12^. The human genome may encode as many as 1000 miRNAs and most genes are subject to regulation by multiple miRNAs^13^. miRNAs have been shown to affect diverse cellular pathways critical to human development and diseases. Accumulating evidence suggests that various miRNAs are aberrantly expressed in cancer cells, including CRC^14^, underscoring the importance of elucidating the mechanism by which miRNAs recognize and regulate their targets.

The advancement of next-generation sequencing techniques has made it possible to generate and analyze vast amounts of high-throughput genomic data, including gene expression profiles and RNA sequencing data^15,16^. However, experimental detection and investigation of miRNA targets and miRNA-induced changes on cellular function is challenging due to the large number of potential interactions to be examined, which is expensive and time-consuming^17–20^. Leveraging computational approaches to predict the potential targets of miRNAs simplifies the process, enabling an initial selection to decrease the number of target sites requiring experimental validation. The earliest computational methods for target prediction mostly employ expert-based knowledge to categorize miRNA-mRNA pairs^21–25^. These methods heavily rely on pre-designed features that have been shown to influence miRNA-mRNA interactions, and the underlying intrinsic mechanisms of the binding process remain incompletely elucidated. Additionally, the necessity of calculating interaction metrics based on sequence data often introduces a laborious computation cost and extra burden to the process, subsequently elongating the execution time for inference. With the development of AI techniques and an increasing number of experimentally validated miRNA-mRNA pairs, many classic machine learning methods have been applied to miRNA target prediction, including support vector machine^26,27^, Naïve Bayes^28,29^ and neural networks^30–33^. For example, Yousef et al. described a target prediction named NBmiRTar^28^ using a naïve Bayes classifier through sequence and miRNA-mRNA duplex information from validated targets and artificially generated negative examples. Lee et al. presented deepTarget^31^, an end-to-end learning framework using deep recurrent neural networks for miRNA target prediction without the need for manual feature extraction. Wen et al. developed DeepMirTar^32^, a stacked de-noising auto-encoder, that combined expert-designed features, e.g., seed match, free energy, sequence composition, and raw sequence data to predict human miRNA targets. These models can extract and learn the feature representations from miRNA-mRNA pairs, predicting the likelihood of binding and improving the models’ performance and efficiency compared with expert-based approaches.

Moreover, the advent of graph neural networks (GNNs)^34–36^ has recently gained significant attention and been applied to a variety of bioinformatic problems such as protein-protein interaction^37,38^, RNA-disease association identification^39,40^, RNA subcellular localization prediction^41,42^, as well as RNA-RNA association prediction^43,44^. Since miRNA-induced silencing complex (miRISC) molecules directly attach to the targeted RNAs, creating intricate graph-like and spatial secondary structures, GNNs present great potential to identify RNA-RNA associations in an end-to-end manner through graph representation of the duplex that can better learn complex interactions between RNAs in a regulatory network. He et al. presented a graph convolutional neural network approach for predicting circRNA-miRNA interactions^45^. Zhao et al. proposed a semantic embedded bipartite graph network for predicting long noncoding RNA-miRNA associations with a novel feature extraction method by combining segmentation, Gaussian interaction profile and graph convolution network^43^. Wang et al. designed a sequence pre-training-based graph neural network to predict lncRNA–miRNA associations from RNA sequences by converting the existing interactions represented as a graph^44^. However, GNN has not been effectively applied to miRNA-mRNA target identification in cancers, specifically in CRC.

To leverage the power of next-generation sequencing techniques and graph-based representations, in this paper, we developed a GNN-based framework using only RNA sequences extracted from CRC cell line (HCT116) to identify potential miRNA targets. We first generate experimental miRNA-mRNA interaction pairs based on AGO-CLASH (UV crosslinking and sequencing of hybrids) method^46^. We then created two pre-trained models to calculate the distributed representations of *k*-mer for input miRNA and mRNA sequences to extract attribute characteristics. We transformed these *k*-mers into nodes for generating node features and graph construction. We finally fed the encoded attribute features of nodes into graph neural networks to detect miRNA-mRNA interactions in CRC. The overall architecture of the proposed graph-based framework is presented in **Figure 1**. The experimental results indicate that our framework could uncover hidden associations between miRNAs and mRNAs, efficiently and accurately identifying miRNA biomarkers that can be used for therapeutic targets in CRC. In the end, we compared our proposed framework to several state-of-the-art methods and demonstrated the superiority of our model in identifying miRNA targets in CRC.

**Figure 1.**
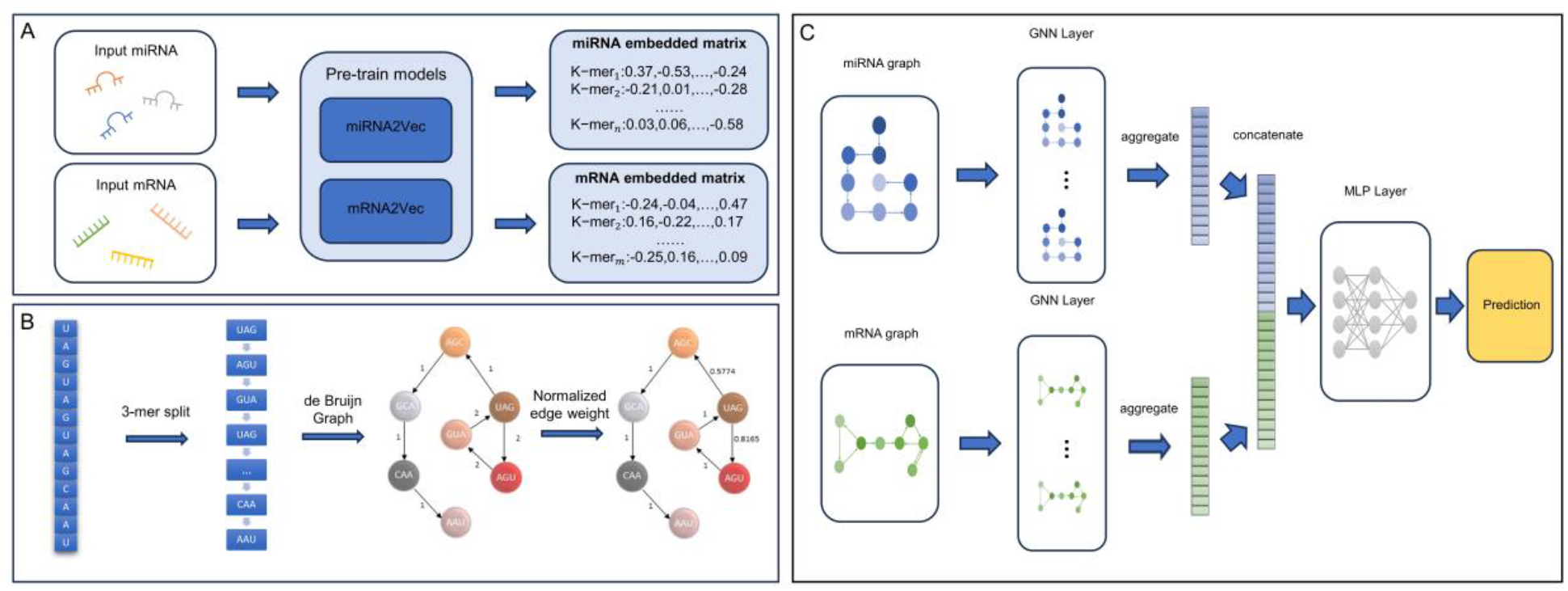
The overall architecture of the proposed framework Gra-CRC-miRTar. (**A**) The creation of pre-training models miRNA2Vec and mRNA2Vec. (**B**) Semantic node features extraction and graph construction. We use 3-mer as an example for the splitting and embedding of the RNA sequences. (**C**) Feature integration with graph neural networks for miRNA target prediction.

## Materials and methods

### Datasets

To obtain experimentally verified miRNA-mRNA interaction pairs in CRC, we utilized AGO-CLASH^47^ data from colorectal cancer (CRC) HCT116 cells, accessible through the NCBI database^48^ (GSE164634). Initially, adapter sequences were removed using Cutadapt software^49^ (version 3.4). Subsequently, the trimmed pair-end FASTQ files were merged employing PEAR software^50^ (version 0.9.6). Each FASTQ file was then collapsed to a single sequence per unique read using Fastx_collapser (version 0.0.14) in the FASTX-Toolkit^51^. Additionally, we trimmed the 5’ and 3’ ends of Unique Molecular Identifiers (UMIs) using Cutadapt^49^ to prepare the sequences for further analysis. To identify interacted miRNA-target hybrids, we analyzed the cleaned FASTA files using Hyb^47^, a bioinformatics pipeline for processing high-throughput cDNA sequencing data from CLASH experiments. To improve the specificity of experimental interaction pairs, we only selected miRNA-mRNA hybrids, and those pairs with minimum interaction energy (ΔG) higher than -11.1 kcal/mol were excluded, as they were deemed indicative of non-specific binding^52^. The remaining pairs were labeled as the positive miRNA-target hybrids. Differently, to obtain negative miRNA-target hybrids, we further processed the cleaned FASTA files by mapping them to a human transcript database using Bowtie2^53^. Gene abundance was calculated based on the totality of mapped reads. The top 100 abundant genes, which were not identified in the positive miRNA-target hybrids were defined as negative controls. Subsequently, reads corresponding to these negative control genes were extracted from the SAM files. Finally, reads from these negative transcripts were randomly connected to miRNAs listed in the positive miRNA-target hybrids to form negative miRNA-target hybrids.

The datasets to construct pre-training models for miRNA and mRNA sequences were obtained from RNAcentral miRbase^54^ and Ensembl^55^ databases, respectively. RNAcentral^56^ is a comprehensive non-coding RNA (ncRNA) sequence collection representing all ncRNA types from a wide variety of organisms and 51 Expert Databases, with over 34 million RNA sequences in different categories. We extracted raw miRNA sequences from the top 29 mammalian species with the largest miRNA size including humans, mice and hamsters, etc. We removed the duplicates of selected species and ended up with 17,275 unique miRNA sequences ranging from 15 nt to 30 nt in length, which will be utilized as miRNA corpus to create the miRNA pre-trained model. In alignment, we selected the same host species of miRNA to collect mRNA sequences from the Ensembl database. We filtered other RNA categories and only kept mRNA sequences from the collection. The average length of the remaining mRNA sequences is 2647.14, where the longest mRNA is 123,179 nt and the shortest is 35 nt. A total of 1,090,566 mRNA sequences were obtained as mRNA corpus after duplicate removal for the construction of the mRNA pre-trained model. The distribution and characteristics of RNA corpus collection on each host species can be found in Supplementary Materials S1.

### Pre-trained model

We used *k*-mer methods to explore the semantic features of RNA sequences. Specifically, the *k*-mer units in RNA sequences exhibit similar structures as words in sentences. Therefore, employing continuous distributed word representations of *k*-mer allows for a natural representation of the contextual information of nucleotides in RNA sequences. We segmented the raw RNA sequences into subsequences by sliding windows and the length of this window is *K*, thus, each subsequence is a *k*-mer. For instance, we assume an RNA sequence contains *N* nucleotides, and it will generate *N* - *K* + 1 overlapping subsequences. We then performed unsupervised training on the collected miRNA and mRNA corpus to construct pre-trained models (miRNA2Vec and mRNA2Vec) based on word2vec^57^ that characterize miRNA and mRNA sequences, respectively. We selected Skip-gram in our experiments to predict the context surrounding a given targeted *k*-mer. During the training process, we utilized negative sampling^58^ and softmax^59^ to optimize the update procedure over all words. We finally decomposed the aggregated model by *k*-mer lengths. After training, we can obtain high-quality and relatively low-dimensional vectors to represent *k*-mer subsequences. Here, we set the parameter *k* to 3∼6 to train the RNA dataset and finally get the embedded vectors. We applied fine-tuning strategy to determine the value of *k* with the best predictive performance. **Figure 2** illustrates the process of the semantic pre-training process of RNA sequences.

**Figure 2.**
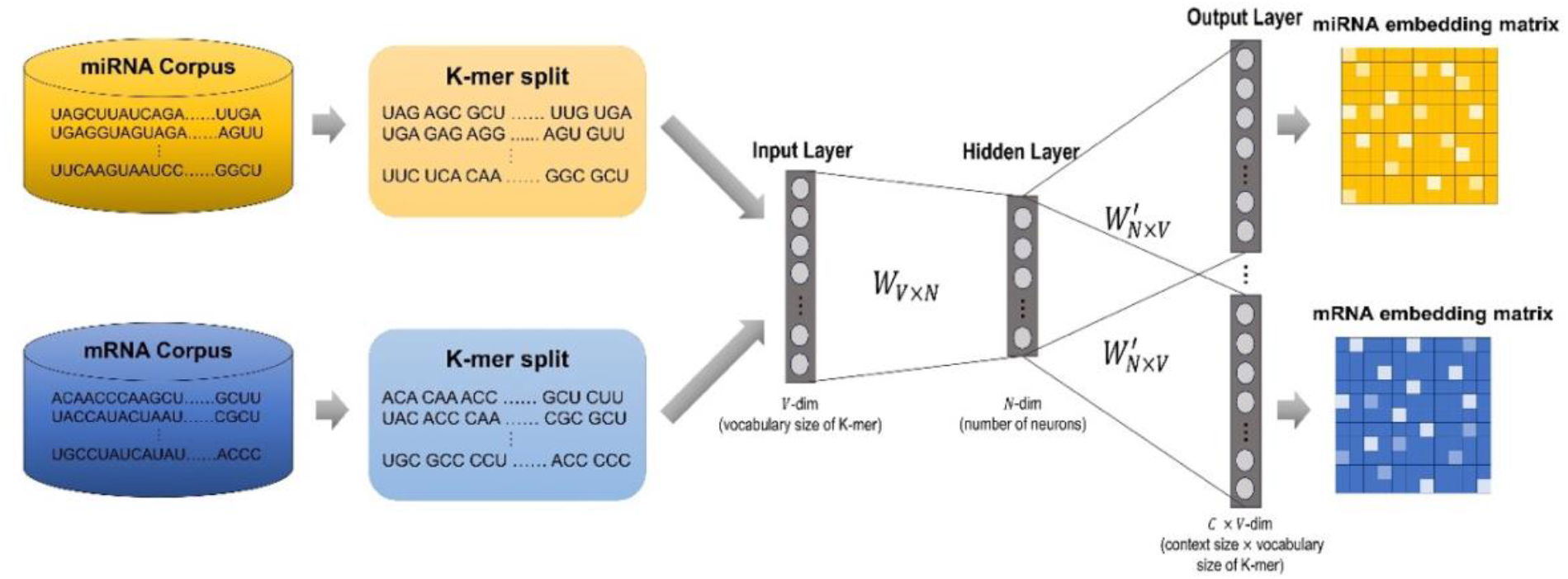
The construction of pre-training of semantic embedding process with Skip-gram architecture.

### Nucleotide to graph

We encoded input RNA sequences into the embedded matrix based on established pre-trained models miRNA2Vec and mRNA2Vec (**Figure 1A**) and transformed miRNA and mRNA sequences into directed graph representation through node features (**Figure 1B**). Specifically, for a given RNA sequence (miRNA or mRNA), we denoted it as (S_1_, S_2_, S_3_, …, S_L-1_, S_L_), where S is one of the nucleotide bases and L is the length of the RNA sequence. For instance, we selected *k*=3 as an example, and the *k*-mer composition set is denoted as {S_1_S_2_S_3_, S_2_S_3_S_4_, …, S_L-2_S_L-1_S_L_}. After *3*-mer segmentation, we assigned these *3*-mers as nodes, following the order of the *3*-mer composition set. We added these nodes one by one to form a de Bruijn graph^60,61^. Subsequently, we allocated weights to each directed edge, with each weight representing the frequency of an edge that connected two 3-mer nodes in the graph. To mitigate the impact of the absolute difference between edge frequencies, we normalized the edge weights in the graph as follows, where *eji* denotes the frequency weight of the edge from node *j* to node *i*, and *N*_*i*_ is the set of neighbor nodes of node *i*.

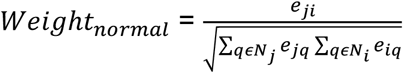

### Graph neural networks

To make full use of the attributes of nodes generated from RNA sequences and improve the feature difference between semantic embedded node features, we leveraged GNNs that can better represent the graph structure characteristics of the nodes. We obtained node features representing *k*-mers of each input RNA sequence after the construction of de Bruijn graph. We then leveraged GNNs to extract and integrate high-level embedded features from the de Bruijn graph (**Figure 1C**). The graph topology and node features generated from mRNA and miRNA sequences were fed into a set of GNN layers, respectively. In this work, we tested and compared three different GNN architectures in Gra-CRC-miRTar, including graph convolutional networks^62^ (GCNs), graph attention networks^63^ (GATs) and graph isomorphism networks^64^ (GINs). The output feature vector of each paired miRNA-mRNA sequence after graph layers were concatenated, followed by a 2-layer MLP for the final prediction.

#### Graph convolutional networks

The GCNs were originally proposed by Kipf and Welling^62^ for semi-supervised learning on graph-structured data based on efficient variants of convolutional neural networks. The propagation rule of GCN is formulated by the following equation to update the network parameters:

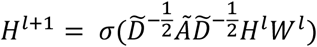

where *Ã* = *A + In* is the adjacency matrix of the graph with added self-connections and *In* is the identify matrix. 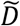 denotes the degree matrix of the adjacency matrix *Ã, W*^*l*^ and *H*^*l*^ indicate the weight and the embedding matrix of the *l*^*th*^ layer, respectively, and *σ* is the non-linear activation function. The fundamental concept behind the GCN layer involves acquiring a transformation function to create a new embedding matrix *H*^*l+*1^ for node *i*. This is implemented by aggregating the intrinsic characteristics of the nodes and the neighboring features of the nodes with normalized edge weights. Through the integration of multiple GCN layers, we can implement inter-node message passing and capture the semantic features of the graph from RNA sequences. More specifically, GCN aggregates the embedded matrixes of all nodes and generates the final graph representation with the readout function, such as mean_pooling, max_pooling and min_pooling, on the learned node representations. Finally, we fed the graph encoding vector to a 2-layer MLP with a ReLU activation function to predict the miRNA targets in CRC.

#### Graph attention networks

Different from GCNs, GATs employ self-attention mechanisms and adapt them to the context of graph data. The fundamental idea behind GATs is to enable nodes in a graph to selectively aggregate information from their neighbors, prioritizing certain nodes or edges over others, based on learned attention coefficients. For each node *i* in the graph, the attention mechanism computes the attention coefficients by considering the features of both the central node and its neighbors. The mathematical equation representing the attention mechanism in GATs is defined as follows:

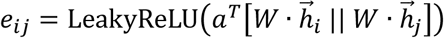

where *eij* represents the unnormalized attention score for the node and *a* is a learned weight vector that is applied to the concatenated node feature representations. LeakyReLU is an activation function that introduces small gradients for negative inputs, and 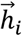 and 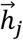 are the feature representations of nodes *i* and *j*, respectively. *W* is the weight matrix applied to the node features and the symbol || denotes the concatenation of the node features. Once *eij* is computed for all node pairs, the scores will go through a Softmax to obtain normalized attention coefficients that sum up to 1 for each node:

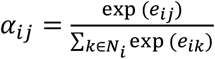

where *αij* is the normalized attention coefficient between node *i* and node *j*, and *Ni* represents the set of neighboring nodes of node *i*. The attention coefficients *αij* will be obtained through Softmax and are used to weigh the representations of neighboring nodes when aggregating information for the central node. This weighted aggregation is carried out for each node in the graph, allowing them to update their own representations based on the information from their neighbors. The node representation is then calculated by summing all neighboring embeddings and the corresponding weights. Similar to the GCN, a readout function is finally applied to obtain the graph representation for the prediction.

#### Graph isomorphism networks

GIN is designed to learn embeddings of graphs that are invariant under graph isomorphism^64^. The main idea of GIN lies in message passing and aggregation mechanisms that iteratively update node representations by considering the local neighborhood structure. These networks aim to capture important graph properties and structural information while being invariant to node permutations. Each GIN involves an aggregation function that aims to iteratively update node representations considering their local neighborhood structure in the graph. It is assumed that 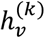 is the hidden representation of node *v* at the *k*-th iteration and the node representations are updated below:

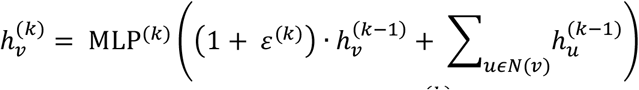

where *N(v)* represents the neighborhood of node *v* and *ε*^*(k)*^ is a parameter that aids in preserving permutation invariance. MLP refers to a multi-layer perceptron that transforms the aggregated information to the sum of the current node representation and the sum of the representations of its neighbors. The use of an MLP helps the GIN to capture complex and nonlinear relationships between nodes and their neighborhoods and enables the learning of invariant graph representations that remain consistent for isomorphic graphs. The aggregation and transformation processes are designed to ensure that the network produces the same output regardless of the ordering or labeling of nodes in isomorphic graphs.

### Experimental setup

#### Implementation and evaluation

We utilized Gensim package^65^ 4.3.0 for the implementation of word2vec embeddings and the construction of pre-trained models. All the models are implemented through Scikit-learn^66^ and PyTorch^67^. For the generated 247,700 paired miRNA-mRNA samples in the dataset, we randomly selected 90% of cases as the training set and 10% as the testing set. We labeled the interacted miRNA-mRNA pairs as “1” and, otherwise, as “0”, which denote positive and negative samples, respectively. We constructed and trained our models using the training set with 5-fold cross-validation and evaluated its capability on the testing set for miRNA target prediction. For GNN-based models, we applied a minimum batch size of 128 for optimization. The learning rate is 0.001 and a drop-out strategy was performed with a rate of 0.3. All the models are iterated for 150 training epochs. For the ablation study, we investigated how different *k*-mers could influence the model’s performance. Additionally, we retrieved 201 experimentally validated CRC-specific miRNA-target pairs from miRTarBase which are not included in our training and testing datasets for external evaluation. We adopted six different metrics to evaluate the predictive performance for all models on the testing set, including accuracy, precision, recall, F1-score, the area under Receiver operating characteristic (AUROC) and the area under Precision-Recall (AUPR) curves.

#### Baseline approaches

We compared our proposed framework with two types of baseline methods for identifying miRNA targets in CRC. The first category involves classic deep learning algorithms, namely, convolutional neural networks (CNNs), recurrent neural networks (RNNs) with the gated recurrent unit (GRU), Bidirectional GRU (BiGRU) and the combinations of these architectures involving attention mechanisms, including CNN + GRU, CNN + BiGRU, GRU + attention mechanism and BiGRU + attention mechanism. The parameters and hyperparameters of these approaches can be found in Supplementary Materials S2. The second category of the baselines is the existing state-of-the-art model developed by others for miRNA target prediction. Here, we selected five representatives, namely, preMLI^68^, CIRNN^69^, LncmirNet^70^, Pmlipred^71^ and PmliHFM^72^. A brief description of these models is shown in Supplementary Materials S3. For the first category baseline approaches, we used the same feature matrix generated from our constructed pre-trained models. For the implementation of others’ models, we followed the original settings as a fair comparison. We also ensured that the training and testing datasets were identical for all compared methods.

## Results

### Comparative performance between our model and classic deep learning classifiers

We compared our proposed framework with several classic architectures of deep neural networks involving both CNN and RNN models, as well as the combination of these models with attention mechanisms. We kept the same experimental setup, such as training and testing set when predicting miRNA targets based on pre-trained embedded features. We used 5-mer as subsequences to construct graph-based models that proved to be the best *k-*mer for the performance (**Table 2**). We conducted the experiments repeatedly five times using different random seeds to split training and testing sets. The prediction outcomes were averaged as shown in **Table 1** with standard deviation in the bracket, from which we can observe that all the models obtained decent predictive performance. In more detail, we found that our proposed framework presents comparable results compared with other deep learning models. Gra-CRC-miRTar with GIN architecture achieved the best values in accuracy (0.887), precision (0.881), F1-score (0.888), AUROC (0.958), and AUPR (0.969), while GRU model with attention mechanisms show better recall values, respectively. Even though our proposed model with GCN and GAT architectures did not display superior performance as with GIN, it still outperformed several other baseline models, such as CNN and CNN+GRU. This is probably because our task is a graph classification problem since we transformed RNA sequences into graphs. We know that GIN is designed to be maximally expressive in the sense of the Weisfeiler-Lehman graph isomorphism test, which allows GIN to better capture the local graph structures up to isomorphism and distinguish between different graph topologies more effectively than GCN and GAT^64^. Meanwhile, comparing the mean aggregator in GCN and the attention mechanisms in GAT, using the sum-based aggregator in GIN to update the node representations enables GNN to capture more discriminative features about the graph.

**Table 1.**
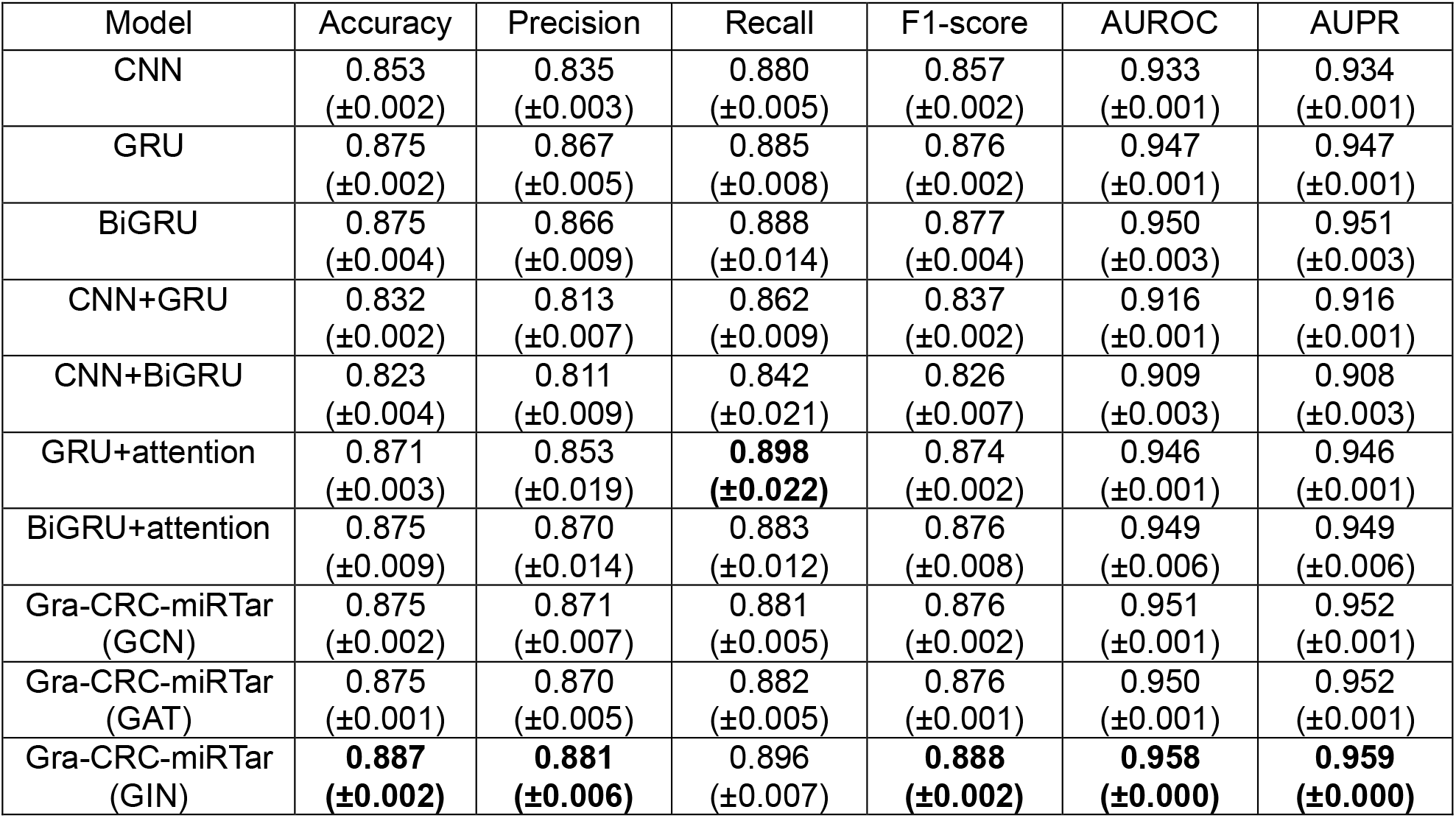
Comparison between our proposed framework with three different GNN architectures and baseline deep learning classifiers.

| Model | Accuracy | Precision | Recall | F1-score | AUROC | AUPR |
| --- | --- | --- | --- | --- | --- | --- |
| CNN | 0.853<br>($\pm 0.002$) | 0.835<br>($\pm 0.003$) | 0.880<br>($\pm 0.005$) | 0.857<br>($\pm 0.002$) | 0.933<br>($\pm 0.001$) | 0.934<br>($\pm 0.001$) |
| GRU | 0.875<br>($\pm 0.002$) | 0.867<br>($\pm 0.005$) | 0.885<br>($\pm 0.008$) | 0.876<br>($\pm 0.002$) | 0.947<br>($\pm 0.001$) | 0.947<br>($\pm 0.001$) |
| BiGRU | 0.875<br>($\pm 0.004$) | 0.866<br>($\pm 0.009$) | 0.888<br>($\pm 0.014$) | 0.877<br>($\pm 0.004$) | 0.950<br>($\pm 0.003$) | 0.951<br>($\pm 0.003$) |
| CNN+GRU | 0.832<br>($\pm 0.002$) | 0.813<br>($\pm 0.007$) | 0.862<br>($\pm 0.009$) | 0.837<br>($\pm 0.002$) | 0.916<br>($\pm 0.001$) | 0.916<br>($\pm 0.001$) |
| CNN+BiGRU | 0.823<br>($\pm 0.004$) | 0.811<br>($\pm 0.009$) | 0.842<br>($\pm 0.021$) | 0.826<br>($\pm 0.007$) | 0.909<br>($\pm 0.003$) | 0.908<br>($\pm 0.003$) |
| GRU+attention | 0.871<br>($\pm 0.003$) | 0.853<br>($\pm 0.019$) | **0.898**<br>**($\pm 0.022$)** | 0.874<br>($\pm 0.002$) | 0.946<br>($\pm 0.001$) | 0.946<br>($\pm 0.001$) |
| BiGRU+attention | 0.875<br>($\pm 0.009$) | 0.870<br>($\pm 0.014$) | 0.883<br>($\pm 0.012$) | 0.876<br>($\pm 0.008$) | 0.949<br>($\pm 0.006$) | 0.949<br>($\pm 0.006$) |
| Gra-CRC-miRTar<br>(GCN) | 0.875<br>($\pm 0.002$) | 0.871<br>($\pm 0.007$) | 0.881<br>($\pm 0.005$) | 0.876<br>($\pm 0.002$) | 0.951<br>($\pm 0.001$) | 0.952<br>($\pm 0.001$) |
| Gra-CRC-miRTar<br>(GAT) | 0.875<br>($\pm 0.001$) | 0.870<br>($\pm 0.005$) | 0.882<br>($\pm 0.005$) | 0.876<br>($\pm 0.001$) | 0.950<br>($\pm 0.001$) | 0.952<br>($\pm 0.001$) |
| Gra-CRC-miRTar<br>(GIN) | **0.887**<br>**($\pm 0.002$)** | **0.881**<br>**($\pm 0.006$)** | 0.896<br>($\pm 0.007$) | **0.888**<br>**($\pm 0.002$)** | **0.958**<br>**($\pm 0.000$)** | **0.959**<br>**($\pm 0.000$)** |

**Table 2.** Performance comparison of our proposed framework Gra-CRC-miRTar with different *k*-mers on three GNN architectures for miRNA targets prediction in CRC.

| $k$-mer | Gra-CRC-miRTar | Accuracy | Precision | Recall | F1-score | AUROC | AUPR |
| --- | --- | --- | --- | --- | --- | --- | --- |
| 3-mer | GCN | 0.836 | 0.818 | 0.864 | 0.840 | 0.921 | 0.925 |
|  | GAT | 0.826 | 0.805 | 0.859 | 0.831 | 0.912 | 0.917 |
|  | GIN | 0.856 | 0.845 | 0.872 | 0.858 | 0.936 | 0.939 |
| 4-mer | GCN | 0.867 | 0.859 | 0.879 | 0.869 | 0.945 | 0.947 |
|  | GAT | 0.861 | 0.849 | 0.878 | 0.863 | 0.940 | 0.943 |
|  | GIN | 0.881 | 0.876 | 0.888 | **0.882** | 0.954 | 0.956 |
| 5-mer | GCN | 0.875 | 0.871 | 0.882 | 0.876 | 0.951 | 0.952 |
|  | GAT | 0.875 | 0.870 | 0.882 | 0.876 | 0.950 | 0.952 |
|  | GIN | **0.887** | **0.880** | **0.896** | 0.879 | **0.958** | **0.959** |
| 6-mer | GCN | 0.871 | 0.865 | 0.879 | 0.872 | 0.947 | 0.948 |
|  | GAT | 0.872 | 0.868 | 0.878 | 0.873 | 0.948 | 0.949 |
|  | GIN | 0.878 | 0.873 | 0.886 | 0.879 | 0.951 | 0.952 |

### Ablation study

Previous studies have indicated that *k*-mer frequency is one of the most critical parameters that will generate distinct graph representations and, as a result, affect the model performance.

Therefore, we examined different values of *k* in the *k*-mer in our proposed framework for three distinct GNN architectures. We kept the other parameters and hyperparameters the same as in **Table 1** for different *k*-mers. The results indicate that when we selected 5-mer or 6-mer, the models obtained slightly better performance than 3-mer and 4-mer on average (**Table 2**). In each *k*-mer, where *k ∈* {3, 4, 5, 6}, GIN shows better results than GCN and GAT in all the metrics. Among all the combinations of *k*-mer and GNN architectures, we found that GIN with 5-mer demonstrated the best performance in accuracy (0.887), precision (0.880), recall (0.896), AUROC (0.958) and AUPR (0.959) for predicting miRNA targets in CRC, while GIN with 4-mer obtained a better F1-score (0.882). Nevertheless, we observed that there is no significant difference in performance using distinct *k*-mers for the prediction. We finally chose the GIN structure with 5-mer as representative for miRNA target identification in comparison with other state-of-the-art methods for external validation.

### Comparison with other existing tools on miRNA targets identification

To further evaluate the capability of Gra-CRC-miRTar in identifying miRNA targets in CRC, we compared it with several well-known tools on the independent testing set. To make a fair comparison, we re-implemented these models and trained them with the same dataset following the raw settings. We used AUROC and AUPR as metrics, which is the quality measure of binary classification, and the results are shown in **Figure 3**, We could observe that Gra-CRC-miRTar (GIN) demonstrates the best AUROC (0.958) and AUPR (0.960) over other tools. Though Gra-CRC-miRTar (GCN) and Gra-CRC-miRTar (GAT) did not obviously show better performance than preMLI, they significantly exceeded LncMirNet and PmliHFM, and are slightly better than PmliPred and CIRNN for miRNA target prediction in CRC. These results suggest that our proposed framework is an effective tool for predicting miRNA targets in CRC and Gra-CRC-miRTar (GIN) proves to be the best architecture among all the models.

**Figure 3.**
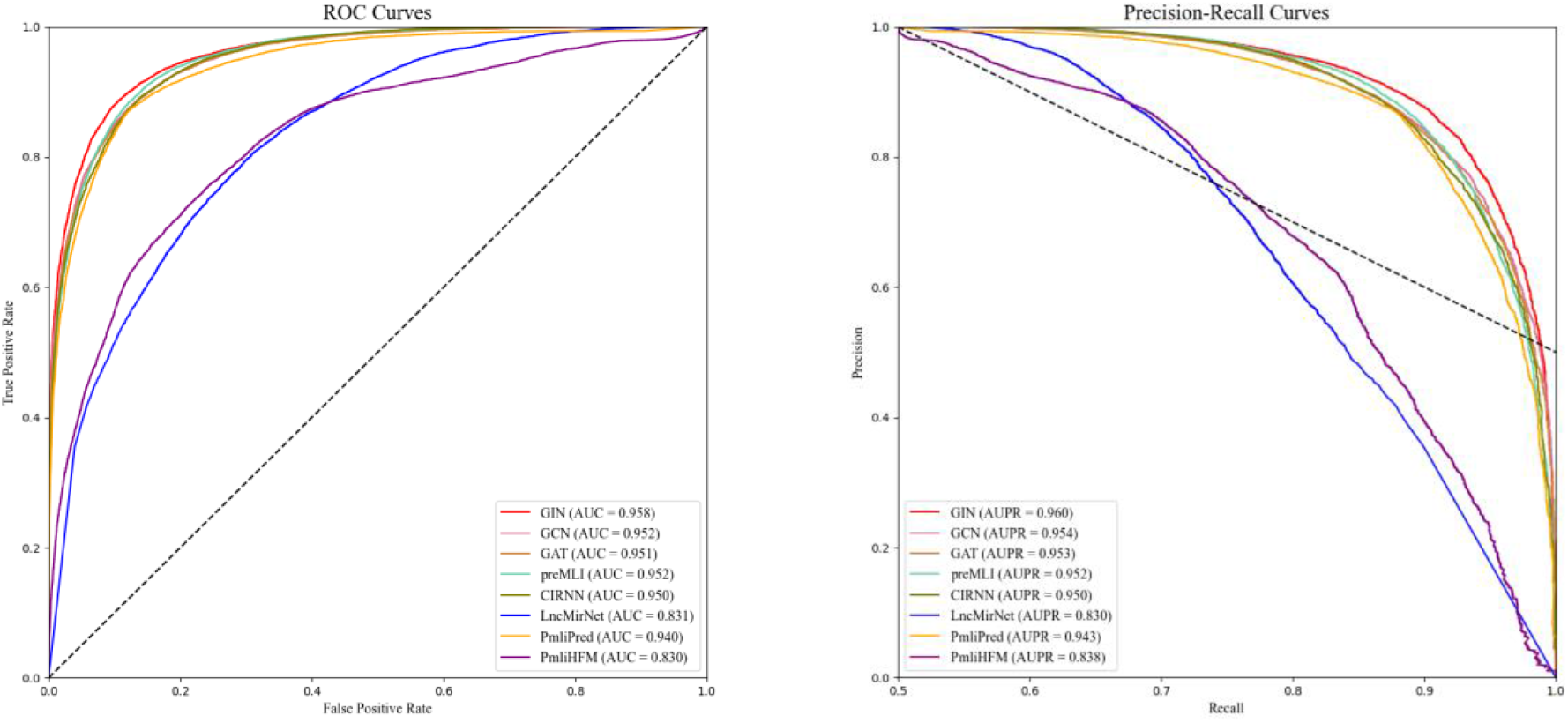
The comparison of ROC and PR curves of Gra-CRC-miRTar and existing state-of-the-art methods for predicting miRNA targets in CRC on the testing set.

### t-SNE visualization of graph vectors

To demonstrate the effectiveness of graph neural networks, we visualized the embedding spaces of feature representations of input RNA sequences before and after the graph-based architectures by projecting them into two dimensions using the t-distributed stochastic neighbor embedding^73,74^ (t-SNE). Figure 4 displays t-SNE visualizations comparing miRNA-mRNA pair feature representations using different *k*-mers, before and after GNN. Here, we selected GIN as the architecture of GNN. The interactive miRNA-mRNA pairs are colored in orange, while the non-interactive ones are in blue. According to the plots, we can find that the two classes before feeding into the graph layers are loosely distributed regardless of the value of *k*. However, we noted that the samples of the same class are separated into clusters after the transformation of GNNs. It is obvious that after learning through the GIN layers, the feature vectors can clearly distinguish between interactive and non-interactive miRNA-mRNA pairs. Interestingly, the visualization shows that the samples of interactive and non-interactive miRNA-mRNA pairs were much more disordered and interweaved before the GNN layer when we used 6-mer for embedding, while we can still gain comparative prediction performance, which further highlights the effectiveness of GNNs for classifying miRNA-mRNA interaction pairs.

**Figure 4.**
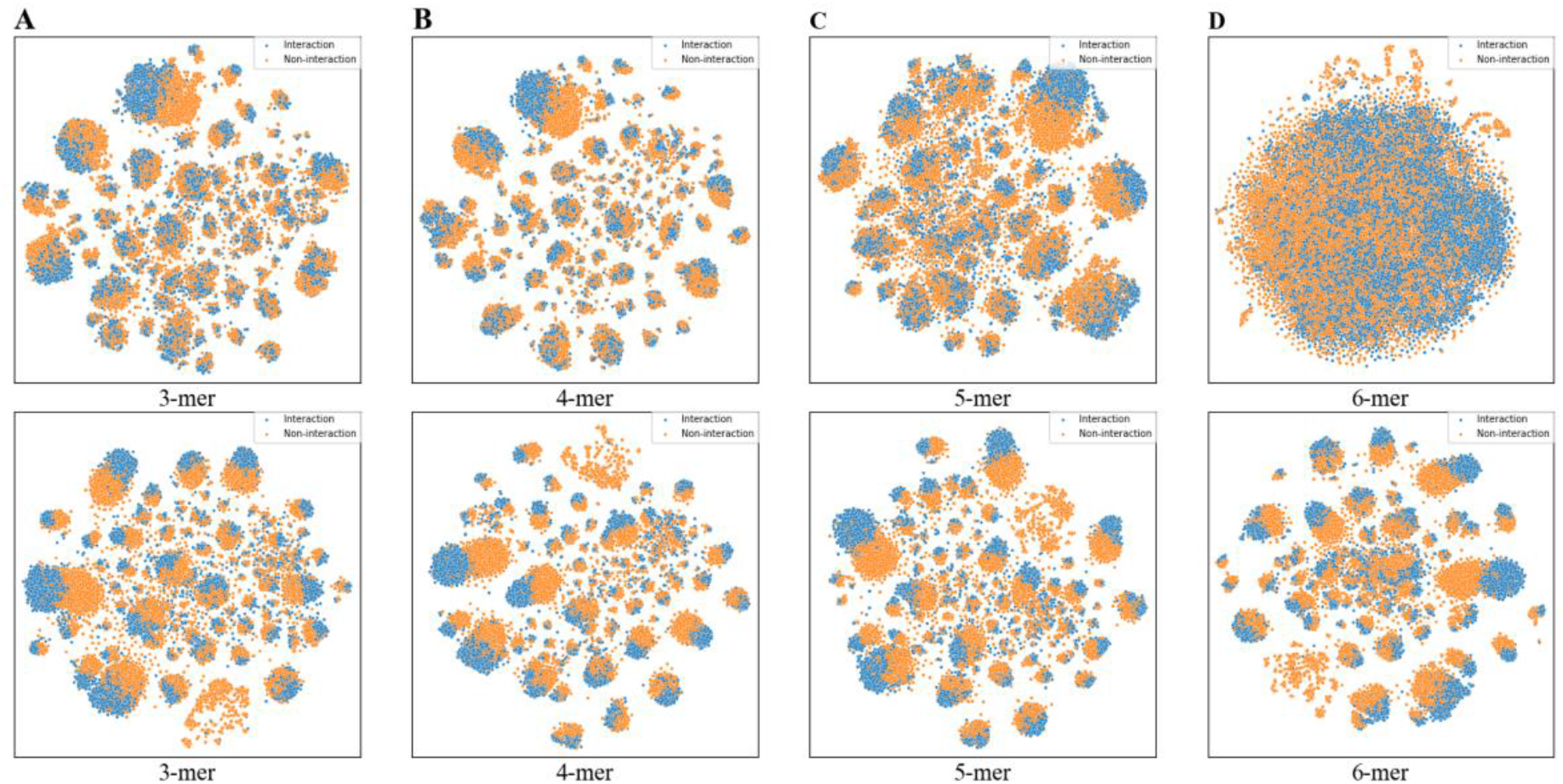
T-SNE visualization of feature vectors of miRNA-mRNA pairs before (top) and after (bottom) GNN based on our constructed pre-trained miRNA2Vec and mRNA2Vec models. Each dot represents a miRNA-target pair, and its color represents its interaction status. (A) k = 3, (B) k = 4, (C) k = 5 and (D) k = 6.

### Novel CRC-specific miRNA target identification through external dataset

To further validate the power of our proposed model, we applied it to an external dataset that contains experimentally validated CRC-specific miRNA–mRNA interactions. We collected 201 new wet lab validated miRNA-target pairs for CRC from miRTarBase^75^ that consist of 75 unique miRNAs and 89 mRNAs. We applied our proposed framework to this dataset along with five other existing methods, including preMLI, CIRNN, LncmirNet, Pmlipred and PmliHFM, to evaluate the miRNA target prediction in CRC. Since this dataset only contains validated interactive pairs, we used recall to measure the performance. The results indicate that our proposed framework can identify 150 (GCN), 146 (GAT), and 158 (GIN) miRNA-target pairs with 0.746, 0.726 and 0.786 in recall, respectively. However, the best tool in comparison is PreMLI obtained 0.716 in recall, followed by LncMirNet (0.701), CIRNN (0.647), PmliPred (0.647) and PmliHFM (0.577). **Table 3** shows the predicted results of 25 miRNA-target pairs with concrete miRNAs and targeted genes for CRC by eight different methods. The enhancement observed could be attributed to the integration of large-scale datasets that our model trained as well as the graph-based methods we selected, which could better capture the underlying patterns of identifying miRNA–target interactions.

**Table 3.** The prediction results on externally validated samples by eight compared methods for miRNA-target identification in CRC. Only 25 out of 201 samples are shown in this table. The complete list of prediction outcomes can be found in Supplementary Materials S4.

| miRTarBaseID | miRNA | Target | CIRNN | Pmlipred | PmliHFM | LncMirNet | PreMLI | GCN | GAT | GIN |
| --- | --- | --- | --- | --- | --- | --- | --- | --- | --- | --- |
| MIRT001190 | hsa-miR-21-5p | PTEN |  |  |  |  | ✓ | ✓ | ✓ | ✓ |
| MIRT003054 | hsa-miR-21-5p | PDCD4 |  | ✓ |  |  | ✓ | ✓ | ✓ | ✓ |
| MIRT003542 | hsa-miR-133a-3p | FSCN1 | ✓ | ✓ | ✓ |  | ✓ | ✓ |  | ✓ |
| MIRT003543 | hsa-miR-145-5p | FSCN1 | ✓ | ✓ | ✓ | ✓ | ✓ | ✓ | ✓ | ✓ |
| MIRT004036 | hsa-miR-185-5p | RHOA | ✓ | ✓ | ✓ | ✓ | ✓ | ✓ | ✓ | ✓ |
| MIRT004037 | hsa-miR-185-5p | CDC42 | ✓ | ✓ | ✓ | ✓ | ✓ | ✓ | ✓ | ✓ |
| MIRT004821 | hsa-miR-34a-5p | E2F1 | ✓ | ✓ | ✓ | ✓ | ✓ | ✓ | ✓ | ✓ |
| MIRT005347 | hsa-miR-103a-3p | DICER1 | ✓ | ✓ | ✓ | ✓ | ✓ | ✓ | ✓ | ✓ |
| MIRT005429 | hsa-miR-21-5p | MSH2 |  | ✓ |  |  |  | ✓ | ✓ | ✓ |
| MIRT005430 | hsa-miR-21-5p | MSH6 | ✓ | ✓ | ✓ | ✓ | ✓ | ✓ | ✓ | ✓ |
| MIRT005553 | hsa-miR-96-5p | KRAS | ✓ | ✓ | ✓ |  | ✓ | ✓ | ✓ | ✓ |
| MIRT005631 | hsa-miR-20a-5p | SMAD4 | ✓ | ✓ | ✓ | ✓ | ✓ | ✓ | ✓ | ✓ |
| MIRT005852 | hsa-miR-17-5p | RBL2 | ✓ | ✓ | ✓ | ✓ | ✓ | ✓ | ✓ | ✓ |
| MIRT005865 | hsa-miR-106b-5p | PTEN |  |  |  | ✓ |  | ✓ | ✓ | ✓ |
| MIRT005869 | hsa-miR-144-3p | NOTCH1 | ✓ | ✓ | ✓ |  | ✓ | ✓ | ✓ | ✓ |
| MIRT682763 | hsa-miR-425-5p | MDM2 | ✓ | ✓ | ✓ | ✓ | ✓ | ✓ | ✓ | ✓ |
| MIRT732287 | hsa-miR-5582-5p | A1BG | ✓ | ✓ |  | ✓ | ✓ |  | ✓ | ✓ |
| MIRT732233 | hsa-miR-138-5p | CD274 |  | ✓ | ✓ | ✓ |  |  |  |  |
| MIRT732288 | hsa-miR-5582-5p | SHC1 | ✓ | ✓ | ✓ | ✓ | ✓ | ✓ |  | ✓ |
| MIRT732305 | hsa-miR-30a-5p | ITGB3 |  | ✓ | ✓ |  | ✓ | ✓ | ✓ | ✓ |
| MIRT002286 | hsa-miR-200c-3p | ZEB1 | ✓ | ✓ | ✓ | ✓ | ✓ | ✓ | ✓ | ✓ |
| MIRT005965 | hsa-miR-330-3p | CDC42 | ✓ |  |  | ✓ |  | ✓ | ✓ | ✓ |
| MIRT006443 | hsa-miR-342-3p | DNMT1 | ✓ |  | ✓ | ✓ | ✓ | ✓ | ✓ | ✓ |
| MIRT006469 | hsa-miR-143-3p | MACC1 |  |  | ✓ | ✓ | ✓ | ✓ |  |  |
| MIRT006664 | hsa-miR-34a-5p | AXL | ✓ | ✓ | ✓ |  | ✓ | ✓ | ✓ | ✓ |

## Discussion

Numerous studies have highlighted the association of miRNAs with cancers, including CRC^76–79^. Nowadays, computational techniques have allowed for the analysis of miRNA targets on a large scale. Yet, many of these approaches frequently generate large amounts of false positives, potentially misrepresenting actual miRNA–mRNA interactions, particularly in disease contexts. Current prediction tools cannot always make accurate and reliable predictions due to the complexity of miRNA targeting, especially in heterogeneous pathological conditions.

Consequently, there is a rationale and need for creating models tailored to specific diseases to reduce the likelihood of incorrect predictions. Although many studies have shown that miRNA function is tissue-specific, so far, few studies have offered an algorithm to predict miRNA targets for a specific disease. The increasingly available RNA data by next-generation sequencing techniques, alongside advancements in natural language processing and graph neural networks, are paving the way for groundbreaking discoveries in genomics and transcriptomics, offering unprecedented insights into complex biological systems and enhancing our ability to understand, diagnose, and treat a vast array of diseases including cancers.

In this study, we developed a novel miRNA target-prediction framework specific for CRC, which is based on pre-trained nucleotide-to-graph neural networks and uses cancer-specific miRNA-target pairs. The high-quality training data was derived from AGO-CLASH experiments, where the precise binding sites of miRNA-mRNA pairs were verified^46^. This foundation enhances the model’s efficiency, as algorithms powered by data are capable of discerning significant and authentic targeting traits within the data. While many current target prediction algorithms aim to achieve high sensitivity in recognizing true positive interactions, they fall short in identifying disease-specific interactions, leading to a higher rate of false positives overall. Our model specifically identifies miRNA targets in CRC with superior performance compared with other benchmark methods in most of the evaluation metrics. One of the possible explanations is that our constructed graph representation can capture the information on the spatial structure of a miRNA-mRNA duplex, which would be beneficial to the prediction of miRNA targets. Our results also revealed that GIN demonstrated the best architecture among all three GNNs that achieved 0.958 in AUROC when using 5-mer for node embedding of RNA sequences, whereas the selection of *k*-mer from 3-mer to 6-mer will not have a significant impact on the prediction results.

We further applied our proposed framework to 201 experimentally validated miRNA-mRNA pairs in CRC from miRTarBase based on western blot, reporter assays, real-time polymerase chain reaction, etc. Our framework successfully identified 172 non-overlapped functional interaction pairs in total using three different GNN structures. The maximum number of predicted targets was 26 for miR-21-5p, followed by 8 and 7 for miR-20a-5p and miRNA-145-5p, respectively. All these predictions can be found in Supplementary Materials S4. The evidence shows that miR-21-5p inhibited the Krev interaction trapped protein 1 (KRIT1) in recipient human umbilical vein endothelial cells, leading to the activation of the β-catenin signaling pathway and an increase in its downstream targets, VEGFa and Ccnd1^80^. This process ultimately enhanced angiogenesis and vascular permeability in CRC, indicating that miR-21-5p may be used as a potential new therapeutic target. Similarly, miR-20a-5p enhanced the invasion and metastasis capabilities of CRC cells by inhibiting Smad4 expression, and elevated levels of miR-20a-5p were associated with a worse prognosis for patients with CRC^81^. The miR-145-5p acts as a suppressor of CRC at the early stage, while promoting CRC metastasis at a late stage through regulating AKT signaling evoked epithelial-mesenchymal transition-mediated anoikis^82^. Our results show that our model is sensitive to discovering target mRNAs whose miRNAs are validated to play critical roles in the regulation of CRC progression, which we believe could serve as an efficient tool to uncover novel dysregulated miRNAs and their targets in CRC.

There are several limitations in this study. First, we lacked a gold standard to collect a set of negative samples as it is challenging to verify truly non-interactive pairs with current techniques. Second, it is essential to delve into and understand the representations learned by GNN-based models to reveal the intrinsic characteristics of the miRNA-mRNA duplex. Gaining these insights will enhance our comprehension of miRNA binding mechanisms and improve our knowledge of the biological processes associated with target prediction. Third, though our proposed model has successfully identified potential novel miRNA-target pairs in CRC, it is critical to further validate their physiological interaction and function at protein levels. Future studies would focus on collecting diverse and larger datasets, particularly those encompassing various tissues and cell types, along with a broader spectrum of miRNA-mRNA interactions for generalizability evaluation. We will improve the model’s interpretability by incorporating explainability methods (e.g., Shapley Additive exPlanations^83^ and Local Interpretable Model-Agnostic Explanations^84^). Moreover, we will build a transfer learning-based model to identify miRNA targets in other cancer types using Gra-CRC-miTar.

## Conclusion

In this paper, we presented a novel framework named Gra-CRC-miTar for miRNA target prediction in CRC. We converted miRNA target prediction into a graph classification task. We created two pre-trained models to encode RNA sequences for graph representation based on word2vec techniques, followed by graph neural networks for the prediction task. The extensive experiments and comprehensive comparison with other methods have demonstrated that Gra-CRC-miTar achieved superior performance for miRNA target prediction in CRC. In addition, our proposed framework successfully identified other experimentally verified miRNA targets with high performance in CRC. Our research introduces a novel path for investigating miRNA-mRNA interactions and creating models that are both more precise and more efficient. As a result, we view our proposed framework as a valuable instrument that has potential applications not only in CRC but also in the identification of miRNA targets for various other diseases.

## Supplementary Materials

The codes and supplementary materials are publicly available at: https://github.com/UF-HOBI-Yin-Lab/Gra-CRC-miRTar

## Acknowledgements

This study was partially supported by grants from the University of Florida Health Cancer Center Pilot Grant AI-2023-03, University of Florida Intelligent Clinical Care Center’s AI2Heal Catalyst Grant, Centers for Disease Control and Prevention (1U18DP006512), National Institute of Environmental Health Sciences (R21ES032762), National Institute of General Medical Sciences (R35GM128753) and the NIH National Center for Advancing Translational Sciences (UL1TR001427).

## Reference

1. Sung, H. et al. Global Cancer Statistics 2020: GLOBOCAN Estimates of Incidence and Mortality Worldwide for 36 Cancers in 185 Countries. CA Cancer J. Clin. 71, 209–249 (2021).

2. Siegel, R. L., Miller, K. D., Wagle, N. S. & Jemal, A. Cancer statistics, 2023. CA Cancer J. Clin. 73, 17–48 (2023).

3. Sharma, R. et al. Global, regional, and national burden of colorectal cancer and its risk factors, 1990–2019: a systematic analysis for the Global Burden of Disease Study 2019. The Lancet Gastroenterology & Hepatology 7, 627–647 (2022).

4. Sawicki, T. et al. A Review of Colorectal Cancer in Terms of Epidemiology, Risk Factors, Development, Symptoms and Diagnosis. Cancers 13, (2021).

5. Rawla, P., Sunkara, T. & Barsouk, A. Epidemiology of colorectal cancer: incidence, mortality, survival, and risk factors. Prz Gastroenterol 14, 89–103 (2019).

6. Arnold, M. et al. Global patterns and trends in colorectal cancer incidence and mortality. Gut 66, 683–691 (2017).

7. Keum, N. & Giovannucci, E. Global burden of colorectal cancer: emerging trends, risk factors and prevention strategies. Nat. Rev. Gastroenterol. Hepatol. 16, 713–732 (2019).

8. Siegel, R. L., Wagle, N. S., Cercek, A., Smith, R. A. & Jemal, A. Colorectal cancer statistics, 2023. CA Cancer J. Clin. 73, 233–254 (2023).

9. Leporrier, J. et al. A population-based study of the incidence, management and prognosis of hepatic metastases from colorectal cancer. Br. J. Surg. 93, 465–474 (2006).

10. Ahluwalia, P., Kolhe, R. & Gahlay, G. K. The clinical relevance of gene expression based prognostic signatures in colorectal cancer. Biochim. Biophys. Acta Rev. Cancer 1875, 188513 (2021).

11. Bazzini, A. A., Lee, M. T. & Giraldez, A. J. Ribosome profiling shows that miR-430 reduces translation before causing mRNA decay in zebrafish. Science 336, 233–237 (2012).

12. Djuranovic, S., Nahvi, A. & Green, R. miRNA-mediated gene silencing by translational repression followed by mRNA deadenylation and decay. Science 336, 237–240 (2012).

13. Bartel, D. P. Metazoan MicroRNAs. Cell 173, 20–51 (2018).

14. Drusco, A. & Croce, C. M. MicroRNAs and Cancer: A Long Story for Short RNAs. Adv. Cancer Res. 135, 1–24 (2017).

15. Levy, S. E. & Myers, R. M. Advancements in Next-Generation Sequencing. Annu. Rev. Genomics Hum. Genet. 17, 95–115 (2016).

16. Hu, T., Chitnis, N., Monos, D. & Dinh, A. Next-generation sequencing technologies: An overview. Hum. Immunol. 82, 801–811 (2021).

17. Gusev, Y., Schmittgen, T. D., Lerner, M., Postier, R. & Brackett, D. Computational analysis of biological functions and pathways collectively targeted by co-expressed microRNAs in cancer. BMC Bioinformatics 8 Suppl 7, S16 (2007).

18. Thomas, M., Lieberman, J. & Lal, A. Desperately seeking microRNA targets. Nat. Struct. Mol. Biol. 17, 1169–1174 (2010).

19. Rojo Arias, J. E. & Busskamp, V. Challenges in microRNAs’ targetome prediction and validation. Neural Regeneration Res. 14, 1672–1677 (2019).

20. Riolo, G., Cantara, S., Marzocchi, C. & Ricci, C. miRNA Targets: From Prediction Tools to Experimental Validation. Methods and Protocols 4, 1 (2020).

21. Lewis, B. P., Shih, I.-H., Jones-Rhoades, M. W., Bartel, D. P. & Burge, C. B. Prediction of mammalian microRNA targets. Cell 115, 787–798 (2003).

22. Rehmsmeier, M., Steffen, P., Hochsmann, M. & Giegerich, R. Fast and effective prediction of microRNA/target duplexes. RNA 10, 1507–1517 (2004).

23. Burgler, C. & Macdonald, P. M. Prediction and verification of microRNA targets by MovingTargets, a highly adaptable prediction method. BMC Genomics 6, 88 (2005).

24. Maragkakis, M. et al. DIANA-microT web server: elucidating microRNA functions through target prediction. Nucleic Acids Res. 37, W273–6 (2009).

25. Betel, D., Koppal, A., Agius, P., Sander, C. & Leslie, C. Comprehensive modeling of microRNA targets predicts functional non-conserved and non-canonical sites. Genome Biol. 11, R90 (2010).

26. Bandyopadhyay, S. & Mitra, R. TargetMiner: microRNA target prediction with systematic identification of tissue-specific negative examples. Bioinformatics 25, 2625–2631 (2009).

27. Liu, H., Yue, D., Chen, Y.Gao, S.-J. & Huang, Y. Improving performance of mammalian microRNA target prediction. BMC Bioinformatics 11, 476 (2010).

28. Yousef, M., Jung, S., Kossenkov, A. V., Showe, L. C. & Showe, M. K. Naïve Bayes for microRNA target predictions—machine learning for microRNA targets. Bioinformatics 23, 2987–2992 (2007).

29. Gaidatzis, D., van Nimwegen, E., Hausser, J. & Zavolan, M. Inference of miRNA targets using evolutionary conservation and pathway analysis. BMC Bioinformatics 8, 69 (2007).

30. Shuang Cheng et al. MiRTDL: A Deep Learning Approach for miRNA Target Prediction. IEEE/ACM Trans. Comput. Biol. Bioinform. 13, 1161–1169 (2016).

31. Lee, B., Baek, J., Park, S. & Yoon, S. deepTarget: End-to-end Learning Framework for microRNA Target Prediction using Deep Recurrent Neural Networks. in Proceedings of the 7th ACM International Conference on Bioinformatics, Computational Biology, and Health Informatics 434–442 (Association for Computing Machinery, New York, NY, USA, 2016).

32. Wen, M., Cong, P., Zhang, Z., Lu, H. & Li, T. DeepMirTar: a deep-learning approach for predicting human miRNA targets. Bioinformatics 34, 3781–3787 (2018).

33. Pla, A., Zhong, X. & Rayner, S. miRAW: A deep learning-based approach to predict microRNA targets by analyzing whole microRNA transcripts. PLoS Comput. Biol. 14, e1006185 (2018).

34. Scarselli, F., Gori, M., Tsoi, A. C., Hagenbuchner, M. & Monfardini, G. The graph neural network model. IEEE Trans. Neural Netw. 20, 61–80 (2009).

35. Wu, Z. et al. A Comprehensive Survey on Graph Neural Networks. IEEE Trans Neural Netw Learn Syst 32, 4–24 (2021).

36. Zhang, X.-M., Liang, L., Liu, L. & Tang, M.-J. Graph Neural Networks and Their Current Applications in Bioinformatics. Front. Genet. 12, 690049 (2021).

37. Réau, M., Renaud, N., Xue, L. C. & Bonvin, A. M. J. J. DeepRank-GNN: a graph neural network framework to learn patterns in protein–protein interfaces. Bioinformatics 39, btac759 (2022).

38. Jha, K., Saha, S. & Singh, H. Prediction of protein–protein interaction using graph neural networks. Sci. Rep. 12, 1–12 (2022).

39. Wang, L. & Zhong, C. gGATLDA: lncRNA-disease association prediction based on graph-level graph attention network. BMC Bioinformatics 23, 11 (2022).

40. Niu, M., Zou, Q. & Wang, C. GMNN2CD: identification of circRNA–disease associations based on variational inference and graph Markov neural networks. Bioinformatics 38, 2246–2253 (2022).

41. Li, M. et al. GraphLncLoc: long non-coding RNA subcellular localization prediction using graph convolutional networks based on sequence to graph transformation. Brief. Bioinform. (2022) doi:10.1093/bib/bbac565.

42. Cai, J., Wang, T., Deng, X., Tang, L. & Liu, L. GM-lncLoc: LncRNAs subcellular localization prediction based on graph neural network with meta-learning. BMC Genomics 24, 52 (2023).

43. Zhao, Z.-Y. et al. SEBGLMA: Semantic Embedded Bipartite Graph Network for Predicting lncRNA-miRNA Associations. Int. J. Intell. Syst. 2023, (2023).

44. Wang, Z. et al. Sequence pre-training-based graph neural network for predicting lncRNA-miRNA associations. Brief. Bioinform. (2023) doi:10.1093/bib/bbad317.

45. He, J. et al. GCNCMI: A Graph Convolutional Neural Network Approach for Predicting circRNA-miRNA Interactions. Front. Genet. 13, 959701 (2022).

46. Fields, C. J. et al. Sequencing of Argonaute-bound microRNA/mRNA hybrids reveals regulation of the unfolded protein response by microRNA-320a. PLoS Genet. 17, e1009934 (2021).

47. Travis, A. J., Moody, J., Helwak, A., Tollervey, D. & Kudla, G. Hyb: a bioinformatics pipeline for the analysis of CLASH (crosslinking, ligation and sequencing of hybrids) data. Methods 65, 263–273 (2014).

48. Schoch, C. L. et al. NCBI Taxonomy: a comprehensive update on curation, resources and tools. Database 2020, (2020).

49. Martin, M. Cutadapt removes adapter sequences from high-throughput sequencing reads. EMBnet.journal 17, 10–12 (2011).

50. Zhang, J., Kobert, K., Flouri, T. & Stamatakis, A. PEAR: a fast and accurate Illumina Paired-End reAd mergeR. Bioinformatics 30, 614–620 (2014).

51. Pearson, W. R., Wood, T., Zhang, Z. & Miller, W. Comparison of DNA sequences with protein sequences. Genomics 46, 24–36 (1997).

52. Helwak, A., Kudla, G., Dudnakova, T. & Tollervey, D. Mapping the human miRNA interactome by CLASH reveals frequent noncanonical binding. Cell 153, 654–665 (2013).

53. Langmead, B. & Salzberg, S. L. Fast gapped-read alignment with Bowtie 2. Nat. Methods 9, 357–359 (2012).

54. Kozomara, A., Birgaoanu, M. & Griffiths-Jones, S. miRBase: from microRNA sequences to function. Nucleic Acids Res. 47, D155–D162 (2019).

55. Cunningham, F. et al. Ensembl 2022. Nucleic Acids Res. 50, D988–D995 (2022).

56. Acids research, N. & 2021. RNAcentral 2021: secondary structure integration, improved sequence search and new member databases. Nucleic Acids Res. 49, D212–D220 (2021).

57. Mikolov, T., Chen, K., Corrado, G. & Dean, J. Efficient Estimation of Word Representations in Vector Space. arXiv [cs.CL] (2013).

58. Goldberg, Y. & Levy, O. word2vec Explained: deriving Mikolov et al.’s negative-sampling word-embedding method. arXiv [cs.CL] (2014).

59. Goodfellow, I., Bengio, Y. & Courville, A. Softmax units for multinoulli output distributions. Deep Learning. Preprint at (2018).

60. Compeau, P. E. C., Pevzner, P. A. & Tesler, G. How to apply de Bruijn graphs to genome assembly. Nat. Biotechnol. 29, 987–991 (2011).

61. Chikhi, R., Limasset, A., Jackman, S., Simpson, J. T. & Medvedev, P. On the representation of de Bruijn graphs. J. Comput. Biol. 22, 336–352 (2015).

62. Kipf, T. N. & Welling, M. Semi-Supervised Classification with Graph Convolutional Networks. arXiv [cs.LG] (2016).

63. Velickovic, P. et al. Graph Attention Networks. arXiv [stat.ML] (2017).

64. Xu, K., Hu, W., Leskovec, J. & Jegelka, S. How Powerful are Graph Neural Networks? arXiv [cs.LG] (2018).

65. Rehurek, R. & Sojka, P. Gensim–python framework for vector space modelling. NLP Centre, Faculty of Informatics, Masaryk University (2011).

66. Pedregosa, F., Varoquaux, G. & Gramfort, A. Scikit-learn: Machine learning in Python. the Journal of machine (2011).

67. Paszke, A. et al. Automatic differentiation in PyTorch. (2017).

68. Yu, X., Jiang, L., Jin, S., Zeng, X. & Liu, X. preMLI: a pre-trained method to uncover microRNA–lncRNA potential interactions. Brief. Bioinform. 23, bbab470 (2021).

69. Zhang, P., Meng, J., Luan, Y. & Liu, C. Plant miRNA-lncRNA interaction prediction with the ensemble of CNN and IndRNN. Interdiscip. Sci. 12, 82–89 (2020).

70. Yang, S. et al. LncMirNet: Predicting LncRNA–miRNA Interaction Based on Deep Learning of Ribonucleic Acid Sequences. Molecules 25, 4372 (2020).

71. Kang, Q., Meng, J., Cui, J., Luan, Y. & Chen, M. PmliPred: a method based on hybrid model and fuzzy decision for plant miRNA–lncRNA interaction prediction. Bioinformatics 36, 2986–2992 (2020).

72. Chen, L. & Sun, Z.-L. PmliHFM: Predicting Plant miRNA-lncRNA Interactions with Hybrid Feature Mining Network. Interdiscip. Sci. 15, 44–54 (2023).

73. Hinton, G. E. & Roweis, S. Stochastic neighbor embedding. Adv. Neural Inf. Process. Syst. 15, (2002).

74. van der Maaten, L. Visualizing Data using t-SNE. https://www.jmlr.org/papers/volume9/vandermaaten08a/vandermaaten08a.pdf?fbcl (2008).

75. Huang, H.-Y. et al. miRTarBase 2020: updates to the experimentally validated microRNA– target interaction database. Nucleic Acids Res. 48, D148–D154 (2019).

76. Garzon, R., Calin, G. A. & Croce, C. M. MicroRNAs in Cancer. Annu. Rev. Med. 60, 167–179 (2009).

77. Peng, Y. & Croce, C. M. The role of MicroRNAs in human cancer. Signal Transduct Target Ther 1, 15004 (2016).

78. Bokhari, A. et al. Targeting nonsense-mediated mRNA decay in colorectal cancers with microsatellite instability. Oncogenesis 7, 70 (2018).

79. He, J. et al. Biomarkers (mRNAs and Non-Coding RNAs) for the Diagnosis and Prognosis of Colorectal Cancer – From the Body Fluid to Tissue Level. Front. Oncol. 11, (2021).

80. He, Q. et al. Cancer-secreted exosomal miR-21-5p induces angiogenesis and vascular permeability by targeting KRIT1. Cell Death Dis. 12, 576 (2021).

81. Cheng, D. et al. MicroRNA-20a-5p promotes colorectal cancer invasion and metastasis by downregulating Smad4. Oncotarget 7, 45199–45213 (2016).

82. Cheng, X. et al. mir-145-5p is a suppressor of colorectal cancer at early stage, while promotes colorectal cancer metastasis at late stage through regulating AKT signaling evoked EMT-mediated anoikis. BMC Cancer 22, 1151 (2022).

83. Lundberg, S. & Lee, S.-I. A unified approach to interpreting model predictions. arXiv [cs.AI] (2017).

84. Ribeiro, M. T., Singh, S. & Guestrin, C. “why should I trust you?”: Explaining the predictions of any classifier. arXiv [cs.LG] (2016) doi:10.1145/2939672.2939778.

