## Supplementary for "Gra-CRC-miRTar: The pre-trained nucleotide-to-graph neural networks to identify potential miRNA targets in colorectal cancer": Supplement 2.docx

The hyperparameter setting of CNN:

For convolution layers:

| Convolution layer type | Convolution1D |
| --- | --- |
| input_shape | (30, 128) (For miRNA) / (90,128) (For mRNA) |
| filters | 64 (For both) |
| kernel_size | 4 (For miRNA)/ 8 (For mRNA) |
| padding | Same (For both) |

For max pooling layers:

| Max pooling layer type | MaxPooling1D |
| --- | --- |
| pool_size | 2 (For miRNA) / 5 (For mRNA) |
| strides | 2 (For miRNA) / 5 (For mRNA) |

The hyperparameter setting of GRU:

| Type | Bidirectional |
| --- | --- |
| units | 50 (For both) |
| return_sequences | True |

The hyperparameter setting of attention layer:

| Dimensionality of the output space | 50 |
| --- | --- |

There are two dense layers network for the final classification:

For the first dense layer:

| units | 64 |
| --- | --- |
| kernel_intitializer | glorot_uniform |
| Normalization Type | BatchNormalization |
| Activation function | relu |
| Dropout | 0.5 |

For the second dense layer:

| units | 1 |
| --- | --- |
| Activation function | sigmoid |

For the optimizer:

| Optimizer Type | Adam |
| --- | --- |
| Learning_rate | 1e-4 |
| Loss function | Binary_crossentropy |
