## Supplementary for "Gra-CRC-miRTar: The pre-trained nucleotide-to-graph neural networks to identify potential miRNA targets in colorectal cancer": Supplement 3 extra validation.docx

| miRTarBaseID | miRNA | MRNA Target | CIRNN | PmliPred | PmliHFM | LncMirNet | PreMLI | GCN | GAT | GIN |
| --- | --- | --- | --- | --- | --- | --- | --- | --- | --- | --- |
| MIRT000046* | *hsa-miR-451a* | *MIF* | ✓ |  | ✓ |  | ✓ | ✓ | ✓ | ✓ |
| MIRT000046* | *hsa-miR-451a* | *MIF* | ✓ | ✓ |  | ✓ | ✓ | ✓ | ✓ | ✓ |
| MIRT000046* | *hsa-miR-451a* | *MIF* | ✓ | ✓ | ✓ | ✓ | ✓ | ✓ | ✓ | ✓ |
| MIRT000316* | *hsa-miR-141-3p* | *ZEB2* | ✓ | ✓ | ✓ | ✓ | ✓ | ✓ | ✓ | ✓ |
| MIRT000316* | *hsa-miR-141-3p* | *ZEB2* | ✓ | ✓ | ✓ | ✓ | ✓ | ✓ | ✓ | ✓ |
| MIRT000316* | *hsa-miR-141-3p* | *ZEB2* | ✓ | ✓ | ✓ | ✓ | ✓ | ✓ | ✓ | ✓ |
| MIRT000636 | *hsa-miR-224-5p* | *CDC42* | ✓ | ✓ | ✓ | ✓ | ✓ | ✓ | ✓ | ✓ |
| MIRT001087* | *hsa-miR-96-5p* | *FOXO1* | ✓ | ✓ |  |  | ✓ | ✓ | ✓ | ✓ |
| MIRT001087* | *hsa-miR-96-5p* | *FOXO1* | ✓ | ✓ | ✓ | ✓ | ✓ | ✓ | ✓ | ✓ |
| MIRT001190* | *hsa-miR-21-5p* | *PTEN* |  |  |  | ✓ |  |  |  |  |
| MIRT001190* | *hsa-miR-21-5p* | *PTEN* | ✓ | ✓ | ✓ |  | ✓ | ✓ | ✓ | ✓ |
| MIRT001190* | *hsa-miR-21-5p* | *PTEN* |  | ✓ | ✓ |  |  |  |  |  |
| MIRT001190* | *hsa-miR-21-5p* | *PTEN* |  | ✓ |  |  | ✓ | ✓ | ✓ |  |
| MIRT001190* | *hsa-miR-21-5p* | *PTEN* |  |  |  |  |  | ✓ |  | ✓ |
| MIRT001190* | *hsa-miR-21-5p* | *PTEN* |  |  |  |  | ✓ | ✓ | ✓ | ✓ |
| MIRT001190* | *hsa-miR-21-5p* | *PTEN* | ✓ | ✓ | ✓ |  | ✓ | ✓ | ✓ | ✓ |
| MIRT003054* | *hsa-miR-21-5p* | *PDCD4* | ✓ | ✓ | ✓ |  | ✓ | ✓ | ✓ | ✓ |
| MIRT003054* | *hsa-miR-21-5p* | *PDCD4* | ✓ | ✓ |  | ✓ | ✓ | ✓ | ✓ | ✓ |
| MIRT003054* | *hsa-miR-21-5p* | *PDCD4* | ✓ | ✓ | ✓ | ✓ | ✓ | ✓ | ✓ | ✓ |
| MIRT003054* | *hsa-miR-21-5p* | *PDCD4* | ✓ | ✓ | ✓ | ✓ | ✓ | ✓ | ✓ | ✓ |
| MIRT003054* | *hsa-miR-21-5p* | *PDCD4* |  | ✓ |  |  | ✓ | ✓ | ✓ | ✓ |
| MIRT003054* | *hsa-miR-21-5p* | *PDCD4* | ✓ | ✓ | ✓ |  | ✓ | ✓ | ✓ | ✓ |
| MIRT003054* | *hsa-miR-21-5p* | *PDCD4* |  | ✓ |  | ✓ | ✓ | ✓ | ✓ | ✓ |
| MIRT003054* | *hsa-miR-21-5p* | *PDCD4* | ✓ | ✓ |  |  | ✓ | ✓ | ✓ | ✓ |
| MIRT003054* | *hsa-miR-21-5p* | *PDCD4* |  | ✓ | ✓ |  | ✓ | ✓ | ✓ | ✓ |
| MIRT003054* | *hsa-miR-21-5p* | *PDCD4* | ✓ |  |  | ✓ |  |  |  |  |
| MIRT003054* | *hsa-miR-21-5p* | *PDCD4* | ✓ |  |  |  | ✓ |  |  |  |
| MIRT003054* | *hsa-miR-21-5p* | *PDCD4* | ✓ | ✓ | ✓ | ✓ | ✓ | ✓ | ✓ | ✓ |
| MIRT003542* | *hsa-miR-133a-3p* | *FSCN1* | ✓ | ✓ | ✓ |  | ✓ | ✓ |  | ✓ |
| MIRT003542* | *hsa-miR-133a-3p* | *FSCN1* | ✓ | ✓ |  | ✓ | ✓ |  |  | ✓ |
| MIRT003542* | *hsa-miR-133a-3p* | *FSCN1* |  |  |  | ✓ |  |  |  | ✓ |
| MIRT003542* | *hsa-miR-133a-3p* | *FSCN1* |  |  |  | ✓ |  |  |  |  |
| MIRT003543* | *hsa-miR-145-5p* | *FSCN1* | ✓ | ✓ | ✓ | ✓ | ✓ | ✓ | ✓ | ✓ |
| MIRT003543* | *hsa-miR-145-5p* | *FSCN1* |  |  | ✓ | ✓ | ✓ | ✓ |  | ✓ |
| MIRT003543* | *hsa-miR-145-5p* | *FSCN1* |  |  |  | ✓ | ✓ | ✓ | ✓ | ✓ |
| MIRT003543* | *hsa-miR-145-5p* | *FSCN1* |  |  | ✓ | ✓ |  | ✓ |  |  |
| MIRT003543* | *hsa-miR-145-5p* | *FSCN1* |  | ✓ |  | ✓ |  |  |  |  |
| MIRT003543* | *hsa-miR-145-5p* | *FSCN1* |  | ✓ |  |  |  |  | ✓ |  |
| MIRT004036* | *hsa-miR-185-5p* | *RHOA* | ✓ | ✓ | ✓ | ✓ | ✓ | ✓ | ✓ | ✓ |
| MIRT004036* | *hsa-miR-185-5p* | *RHOA* | ✓ | ✓ | ✓ | ✓ | ✓ | ✓ | ✓ | ✓ |
| MIRT004037 | *hsa-miR-185-5p* | *CDC42* | ✓ | ✓ | ✓ | ✓ | ✓ | ✓ | ✓ | ✓ |
| MIRT004821 | *hsa-miR-34a-5p* | *E2F1* | ✓ | ✓ | ✓ | ✓ | ✓ | ✓ | ✓ | ✓ |
| MIRT005347* | *hsa-miR-103a-3p* | *DICER1* | ✓ | ✓ | ✓ | ✓ | ✓ | ✓ | ✓ | ✓ |
| MIRT005347* | *hsa-miR-103a-3p* | *DICER1* |  | ✓ |  | ✓ | ✓ | ✓ | ✓ | ✓ |
| MIRT005429* | *hsa-miR-21-5p* | *MSH2* |  | ✓ |  |  |  | ✓ | ✓ | ✓ |
| MIRT005429* | *hsa-miR-21-5p* | *MSH2* | ✓ | ✓ | ✓ | ✓ | ✓ | ✓ | ✓ | ✓ |
| MIRT005429* | *hsa-miR-21-5p* | *MSH2* |  |  | ✓ |  | ✓ | ✓ | ✓ | ✓ |
| MIRT005430 | *hsa-miR-21-5p* | *MSH6* | ✓ | ✓ | ✓ | ✓ | ✓ | ✓ | ✓ | ✓ |
| MIRT005450 | *hsa-miR-96-5p* | *FOXO3* | ✓ |  | ✓ |  | ✓ | ✓ |  |  |
| MIRT005553* | *hsa-miR-96-5p* | *KRAS* | ✓ | ✓ | ✓ |  | ✓ | ✓ | ✓ | ✓ |
| MIRT005553* | *hsa-miR-96-5p* | *KRAS* | ✓ | ✓ | ✓ | ✓ | ✓ | ✓ | ✓ | ✓ |
| MIRT005631* | *hsa-miR-20a-5p* | *SMAD4* | ✓ | ✓ | ✓ | ✓ | ✓ | ✓ | ✓ | ✓ |
| MIRT005631* | *hsa-miR-20a-5p* | *SMAD4* | ✓ |  |  |  | ✓ | ✓ | ✓ | ✓ |
| MIRT005631* | *hsa-miR-20a-5p* | *SMAD4* |  |  |  | ✓ |  | ✓ | ✓ | ✓ |
| MIRT005852 | *hsa-miR-17-5p* | *RBL2* | ✓ | ✓ | ✓ | ✓ | ✓ | ✓ | ✓ | ✓ |
| MIRT005865* | *hsa-miR-106b-5p* | *PTEN* |  |  |  | ✓ |  | ✓ | ✓ | ✓ |
| MIRT005865* | *hsa-miR-106b-5p* | *PTEN* |  |  |  |  |  |  |  |  |
| MIRT005869 | *hsa-miR-144-3p* | *NOTCH1* | ✓ | ✓ | ✓ |  | ✓ | ✓ | ✓ | ✓ |
| MIRT006148* | *hsa-miR-92a-3p* | *BCL2L11* |  |  | ✓ | ✓ | ✓ | ✓ | ✓ | ✓ |
| MIRT006148* | *hsa-miR-92a-3p* | *BCL2L11* |  |  |  | ✓ | ✓ | ✓ | ✓ | ✓ |
| MIRT006148* | *hsa-miR-92a-3p* | *BCL2L11* | ✓ |  | ✓ |  | ✓ | ✓ | ✓ | ✓ |
| miRTarBaseID | *miRNA* | *MRNA Target* | *CIRNN* | *PmliPred* | *PmliHFM* | *LncMirNet* | *PreMLI* | *GCN* | *GAT* | *GIN* |
| MIRT006148* | *hsa-miR-92a-3p* | *BCL2L11* | ✓ |  |  | ✓ | ✓ | ✓ | ✓ | ✓ |
| MIRT006148* | *hsa-miR-92a-3p* | *BCL2L11* | ✓ | ✓ | ✓ |  | ✓ | ✓ | ✓ | ✓ |
| MIRT006199 | *hsa-miR-21-5p* | *CCL20* | ✓ | ✓ |  | ✓ | ✓ | ✓ | ✓ | ✓ |
| MIRT006511* | *hsa-miR-200b-3p* | *RND3* | ✓ | ✓ | ✓ | ✓ | ✓ | ✓ | ✓ | ✓ |
| MIRT006511* | *hsa-miR-200b-3p* | *RND3* | ✓ | ✓ | ✓ | ✓ | ✓ | ✓ | ✓ | ✓ |
| MIRT006511* | *hsa-miR-200b-3p* | *RND3* | ✓ | ✓ | ✓ | ✓ | ✓ | ✓ | ✓ | ✓ |
| MIRT006564* | *hsa-miR-99b-5p* | *MTOR* | ✓ |  |  | ✓ |  |  |  |  |
| MIRT006564* | *hsa-miR-99b-5p* | *MTOR* | ✓ | ✓ | ✓ | ✓ | ✓ | ✓ | ✓ | ✓ |
| MIRT006564* | *hsa-miR-99b-5p* | *MTOR* | ✓ | ✓ | ✓ | ✓ | ✓ | ✓ | ✓ | ✓ |
| MIRT006567* | *hsa-miR-31-5p* | *SATB2* |  | ✓ | ✓ | ✓ | ✓ | ✓ | ✓ | ✓ |
| MIRT006567* | *hsa-miR-31-5p* | *SATB2* |  | ✓ | ✓ | ✓ | ✓ | ✓ | ✓ | ✓ |
| MIRT006567* | *hsa-miR-31-5p* | *SATB2* | ✓ | ✓ | ✓ | ✓ | ✓ | ✓ | ✓ | ✓ |
| MIRT006567* | *hsa-miR-31-5p* | *SATB2* | ✓ | ✓ | ✓ | ✓ | ✓ | ✓ | ✓ | ✓ |
| MIRT006567* | *hsa-miR-31-5p* | *SATB2* |  |  | ✓ |  | ✓ |  |  | ✓ |
| MIRT006663 | *hsa-miR-210-3p* | *VMP1* |  | ✓ |  | ✓ |  | ✓ |  | ✓ |
| MIRT007107 | *hsa-miR-145-5p* | *NRAS* |  |  |  | ✓ | ✓ |  |  | ✓ |
| MIRT024977 | *hsa-miR-204-5p* | *HMGA2* |  |  | ✓ | ✓ |  |  |  |  |
| MIRT036232 | *hsa-miR-320b* | *MYC* | ✓ | ✓ |  | ✓ | ✓ | ✓ | ✓ | ✓ |
| MIRT047724* | *hsa-miR-10a-5p* | *ACTG1* | ✓ | ✓ | ✓ | ✓ | ✓ | ✓ | ✓ | ✓ |
| MIRT047724* | *hsa-miR-10a-5p* | *ACTG1* | ✓ | ✓ | ✓ |  | ✓ | ✓ | ✓ |  |
| MIRT052930 | *hsa-miR-15a-5p* | *REPIN1* |  |  |  |  |  |  |  |  |
| MIRT053250* | *hsa-miR-224-5p* | *PHLPP1* | ✓ | ✓ | ✓ | ✓ | ✓ | ✓ | ✓ | ✓ |
| MIRT053250* | *hsa-miR-224-5p* | *PHLPP1* |  |  |  | ✓ |  | ✓ | ✓ | ✓ |
| MIRT053338* | *hsa-miR-196b-5p* | *FAS* | ✓ | ✓ |  | ✓ | ✓ | ✓ | ✓ | ✓ |
| MIRT053338* | *hsa-miR-196b-5p* | *FAS* | ✓ | ✓ | ✓ |  | ✓ | ✓ | ✓ | ✓ |
| MIRT054372* | *hsa-miR-34a-5p* | *KLF4* |  | ✓ |  | ✓ | ✓ | ✓ | ✓ | ✓ |
| MIRT054372* | *hsa-miR-34a-5p* | *KLF4* |  | ✓ | ✓ |  | ✓ | ✓ | ✓ | ✓ |
| MIRT054425* | *hsa-miR-126-3p* | *CXCR4* | ✓ | ✓ | ✓ | ✓ | ✓ |  | ✓ | ✓ |
| MIRT054425* | *hsa-miR-126-3p* | *CXCR4* | ✓ | ✓ | ✓ | ✓ |  | ✓ | ✓ |  |
| MIRT054440 | *hsa-miR-378a-3p* | *VIM* | ✓ | ✓ |  | ✓ | ✓ | ✓ | ✓ | ✓ |
| MIRT054514 | *hsa-miR-23a-3p* | *APAF1* |  | ✓ | ✓ | ✓ | ✓ | ✓ | ✓ | ✓ |
| MIRT054540 | *hsa-miR-518a-3p* | *CCR6* |  |  | ✓ |  | ✓ | ✓ | ✓ | ✓ |
| MIRT054582* | *hsa-miR-30b-5p* | *SIX1* |  |  |  | ✓ |  |  |  |  |
| MIRT054632* | *hsa-miR-30a-5p* | *HSPA5* | ✓ |  | ✓ |  | ✓ |  |  |  |
| MIRT054632 | *hsa-miR-30a-5p* | *HSPA5* | ✓ | ✓ | ✓ |  | ✓ | ✓ |  |  |
| MIRT054638 | *hsa-miR-214-3p* | *FGFR1* | ✓ | ✓ |  | ✓ |  | ✓ | ✓ | ✓ |
| MIRT054763* | *hsa-miR-139-5p* | *NOTCH1* |  |  | ✓ | ✓ |  | ✓ | ✓ | ✓ |
| MIRT054763* | *hsa-miR-139-5p* | *NOTCH1* | ✓ | ✓ |  | ✓ | ✓ | ✓ | ✓ | ✓ |
| MIRT054763* | *hsa-miR-139-5p* | *NOTCH1* | ✓ | ✓ | ✓ | ✓ | ✓ | ✓ | ✓ | ✓ |
| MIRT069484 | *hsa-miR-543* | *MTA1* | ✓ | ✓ | ✓ | ✓ | ✓ | ✓ | ✓ | ✓ |
| MIRT099904* | *hsa-miR-363-3p* | *SOX4* |  | ✓ | ✓ | ✓ | ✓ | ✓ | ✓ | ✓ |
| MIRT099904* | *hsa-miR-363-3p* | *SOX4* |  |  | ✓ | ✓ | ✓ |  | ✓ | ✓ |
| MIRT099904* | *hsa-miR-363-3p* | *SOX4* |  |  | ✓ | ✓ | ✓ | ✓ | ✓ | ✓ |
| MIRT099904* | *hsa-miR-363-3p* | *SOX4* |  |  | ✓ |  | ✓ |  | ✓ | ✓ |
| MIRT099904* | *hsa-miR-363-3p* | *SOX4* |  |  | ✓ | ✓ | ✓ | ✓ | ✓ | ✓ |
| MIRT099904* | *hsa-miR-363-3p* | *SOX4* |  |  | ✓ | ✓ | ✓ |  | ✓ | ✓ |
| MIRT241527* | *hsa-miR-374a-5p* | *CCND1* | ✓ |  |  |  |  |  |  |  |
| MIRT241527* | *hsa-miR-374a-5p* | *CCND1* | ✓ |  |  | ✓ |  |  |  | ✓ |
| MIRT241527* | *hsa-miR-374a-5p* | *CCND1* | ✓ |  |  |  |  |  |  | ✓ |
| MIRT437392 | *hsa-miR-221-5p* | *MBD2* | ✓ | ✓ | ✓ | ✓ | ✓ |  | ✓ | ✓ |
| MIRT437393 | *hsa-miR-224-5p* | *MBD2* |  |  |  | ✓ |  |  |  |  |
| MIRT437988* | *hsa-miR-200b-3p* | *CDKN1B* |  |  | ✓ |  | ✓ |  |  |  |
| MIRT437988* | *hsa-miR-200b-3p* | *CDKN1B* | ✓ |  | ✓ | ✓ |  | ✓ | ✓ | ✓ |
| MIRT438184 | *hsa-miR-135a-5p* | *SIAH1* |  |  |  | ✓ |  |  |  |  |
| MIRT438195* | *hsa-miR-7-5p* | *PAX6* | ✓ | ✓ |  | ✓ | ✓ | ✓ | ✓ | ✓ |
| MIRT438195* | *hsa-miR-7-5p* | *PAX6* | ✓ | ✓ |  | ✓ | ✓ | ✓ | ✓ | ✓ |
| MIRT438201 | *hsa-miR-138-5p* | *TWIST2* | ✓ | ✓ | ✓ | ✓ |  | ✓ | ✓ | ✓ |
| MIRT438437* | *hsa-miR-181a-5p* | *WIF1* | ✓ | ✓ | ✓ | ✓ | ✓ | ✓ | ✓ | ✓ |
| MIRT438437* | *hsa-miR-181a-5p* | *WIF1* | ✓ | ✓ |  | ✓ | ✓ | ✓ | ✓ | ✓ |
| MIRT438723 | *hsa-miR-100-5p* | *RAP1B* | ✓ | ✓ | ✓ | ✓ | ✓ | ✓ | ✓ | ✓ |
| MIRT513806* | *hsa-miR-302a-3p* | *NFIB* | ✓ | ✓ | ✓ | ✓ | ✓ | ✓ |  | ✓ |
| miRTarBaseID | *miRNA* | *MRNA Target* | *CIRNN* | *PmliPred* | *PmliHFM* | *LncMirNet* | *PreMLI* | *GCN* | *GAT* | *GIN* |
| MIRT513806* | *hsa-miR-302a-3p* | *NFIB* | ✓ | ✓ | ✓ | ✓ | ✓ | ✓ | ✓ | ✓ |
| MIRT557277* | *hsa-miR-543* | *HMGA2* |  | ✓ | ✓ | ✓ | ✓ | ✓ | ✓ | ✓ |
| MIRT557277* | *hsa-miR-543* | *HMGA2* |  |  |  | ✓ |  |  |  |  |
| MIRT682763 | *hsa-miR-425-5p* | *MDM2* | ✓ | ✓ | ✓ | ✓ | ✓ | ✓ | ✓ | ✓ |
| MIRT686573 | *hsa-miR-425-5p* | *TNFRSF10B* | ✓ | ✓ |  |  | ✓ |  | ✓ | ✓ |
| MIRT731556 | *hsa-miR-544a* | *HOXA10* |  | ✓ |  |  | ✓ | ✓ | ✓ | ✓ |
| MIRT731867 | *hsa-miR-146a-5p* | *CPM* |  |  |  |  |  | ✓ | ✓ | ✓ |
| MIRT731978 | *hsa-miR-659-3p* | *SPHK1* | ✓ | ✓ |  | ✓ | ✓ | ✓ | ✓ | ✓ |
| MIRT732177 | *hsa-miR-542-3p* | *CTTN* | ✓ |  |  |  |  |  |  | ✓ |
| MIRT732188 | *hsa-miR-543* | *KRAS* | ✓ | ✓ | ✓ | ✓ |  | ✓ | ✓ | ✓ |
| MIRT732202 | *hsa-miR-106b-5p* | *PRRX1* | ✓ | ✓ | ✓ | ✓ | ✓ | ✓ | ✓ | ✓ |
| MIRT732215 | *hsa-miR-608* | *NAA10* | ✓ | ✓ | ✓ | ✓ | ✓ | ✓ | ✓ | ✓ |
| MIRT732226 | *hsa-miR-490-3p* | *TGFBR1* | ✓ | ✓ |  | ✓ |  | ✓ | ✓ | ✓ |
| MIRT732232 | *hsa-miR-139-5p* | *BCL2* | ✓ | ✓ |  | ✓ | ✓ | ✓ | ✓ | ✓ |
| MIRT732233* | *hsa-miR-138-5p* | *CD274* | ✓ | ✓ | ✓ | ✓ | ✓ |  |  | ✓ |
| MIRT732233* | *hsa-miR-138-5p* | *CD274* |  | ✓ | ✓ | ✓ |  |  |  |  |
| MIRT732238 | *hsa-miR-875-5p* | *EGFR* | ✓ |  |  |  | ✓ |  |  |  |
| MIRT732287 | *hsa-miR-5582-5p* | *A1BG* | ✓ | ✓ |  | ✓ | ✓ |  | ✓ | ✓ |
| MIRT732288 | *hsa-miR-5582-5p* | *SHC1* | ✓ | ✓ | ✓ | ✓ | ✓ | ✓ |  | ✓ |
| MIRT732289 | *hsa-miR-5582-5p* | *CDK2* | ✓ | ✓ |  |  | ✓ | ✓ | ✓ | ✓ |
| MIRT732300 | *hsa-miR-625-3p* | *MAP2K6* | ✓ | ✓ |  |  |  |  |  | ✓ |
| MIRT732303 | *hsa-miR-29a-5p* | *PTEN* | ✓ | ✓ |  |  | ✓ | ✓ | ✓ | ✓ |
| MIRT732305 | *hsa-miR-30a-5p* | *ITGB3* |  | ✓ | ✓ |  | ✓ | ✓ | ✓ | ✓ |
| MIRT732429 | *hsa-miR-384* | *KRAS* |  |  |  | ✓ |  |  |  | ✓ |
| MIRT732430* | *hsa-miR-384* | *FBXW7* |  | ✓ | ✓ | ✓ | ✓ |  |  | ✓ |
| MIRT732430* | *hsa-miR-384* | *FBXW7* |  |  |  |  | ✓ | ✓ |  |  |
| MIRT732956 | *hsa-miR-224-5p* | *CCND1* |  | ✓ |  | ✓ |  |  |  |  |
| MIRT732967 | *hsa-miR-506-3p* | *EZH2* |  |  | ✓ | ✓ |  |  |  |  |
| MIRT733763 | *hsa-miR-135b-5p* | *MUL1* | ✓ |  | ✓ | ✓ |  |  |  | ✓ |
| MIRT733774 | *hsa-miR-488-5p* | *PFKFB3* |  |  |  | ✓ |  |  |  |  |
| MIRT733824 | *hsa-miR-302a-3p* | *CD44* |  | ✓ | ✓ | ✓ | ✓ | ✓ | ✓ | ✓ |
| MIRT733866 | *hsa-miR-484* | *KLF12* |  |  | ✓ | ✓ |  | ✓ | ✓ | ✓ |
| MIRT735431 | *hsa-miR-195-5p* | *YAP1* | ✓ | ✓ | ✓ | ✓ | ✓ | ✓ | ✓ | ✓ |
| MIRT002272* | *hsa-miR-221-3p* | *CDKN1C* |  | ✓ | ✓ | ✓ | ✓ |  | ✓ | ✓ |
| MIRT002272* | *hsa-miR-221-3p* | *CDKN1C* |  | ✓ | ✓ |  | ✓ | ✓ | ✓ | ✓ |
| MIRT002286* | *hsa-miR-200c-3p* | *ZEB1* | ✓ | ✓ | ✓ | ✓ | ✓ | ✓ | ✓ | ✓ |
| MIRT002286* | *hsa-miR-200c-3p* | *ZEB1* | ✓ | ✓ |  |  | ✓ | ✓ | ✓ | ✓ |
| MIRT002286* | *hsa-miR-200c-3p* | *ZEB1* | ✓ |  | ✓ |  | ✓ | ✓ | ✓ | ✓ |
| MIRT002286* | *hsa-miR-200c-3p* | *ZEB1* | ✓ | ✓ | ✓ | ✓ | ✓ | ✓ | ✓ | ✓ |
| MIRT003317* | *hsa-miR-21-5p* | *RHOB* | ✓ | ✓ | ✓ | ✓ | ✓ | ✓ | ✓ | ✓ |
| MIRT003317* | *hsa-miR-21-5p* | *RHOB* | ✓ | ✓ | ✓ |  | ✓ | ✓ | ✓ | ✓ |
| MIRT004285 | *hsa-miR-491-5p* | *BCL2L1* | ✓ | ✓ |  | ✓ |  | ✓ | ✓ |  |
| MIRT004402* | *hsa-miR-34c-5p* | *MYC* | ✓ | ✓ |  | ✓ | ✓ | ✓ | ✓ | ✓ |
| MIRT004402* | *hsa-miR-34c-5p* | *MYC* | ✓ | ✓ |  | ✓ | ✓ | ✓ | ✓ | ✓ |
| MIRT005319* | *hsa-miR-34b-3p* | *MYC* |  |  |  | ✓ | ✓ |  | ✓ | ✓ |
| MIRT005319* | *hsa-miR-34b-3p* | *MYC* |  |  |  | ✓ |  |  |  |  |
| MIRT005481* | *hsa-miR-20a-5p* | *BNIP2* | ✓ |  | ✓ |  |  | ✓ | ✓ | ✓ |
| MIRT005481* | *hsa-miR-20a-5p* | *BNIP2* | ✓ |  |  | ✓ |  | ✓ |  | ✓ |
| MIRT005481* | *hsa-miR-20a-5p* | *BNIP2* | ✓ | ✓ | ✓ | ✓ | ✓ | ✓ | ✓ | ✓ |
| MIRT005481* | *hsa-miR-20a-5p* | *BNIP2* | ✓ |  | ✓ | ✓ | ✓ | ✓ | ✓ | ✓ |
| MIRT005481* | *hsa-miR-20a-5p* | *BNIP2* | ✓ |  |  | ✓ | ✓ |  |  |  |
| MIRT005759 | *hsa-miR-148b-3p* | *CCKBR* | ✓ | ✓ |  | ✓ | ✓ | ✓ | ✓ | ✓ |
| MIRT005965* | *hsa-miR-330-3p* | *CDC42* | ✓ | ✓ | ✓ | ✓ | ✓ | ✓ | ✓ | ✓ |
| MIRT005965* | *hsa-miR-330-3p* | *CDC42* | ✓ |  |  | ✓ |  | ✓ | ✓ | ✓ |
| MIRT006443 | *hsa-miR-342-3p* | *DNMT1* | ✓ |  | ✓ | ✓ | ✓ | ✓ | ✓ | ✓ |
| MIRT006469 | *hsa-miR-143-3p* | *MACC1* |  |  | ✓ | ✓ | ✓ | ✓ |  |  |
| MIRT006664* | *hsa-miR-34a-5p* | *AXL* | ✓ | ✓ | ✓ | ✓ | ✓ | ✓ | ✓ | ✓ |
| MIRT006664* | *hsa-miR-34a-5p* | *AXL* | ✓ | ✓ | ✓ |  | ✓ | ✓ | ✓ | ✓ |
| MIRT006688* | *hsa-miR-103a-3p* | *KLF4* | ✓ | ✓ | ✓ | ✓ | ✓ | ✓ | ✓ | ✓ |
| miRTarBaseID | *miRNA* | *MRNA Target* | *CIRNN* | *PmliPred* | *PmliHFM* | *LncMirNet* | *PreMLI* | *GCN* | *GAT* | *GIN* |
| MIRT006688* | *hsa-miR-103a-3p* | *KLF4* |  |  |  |  |  |  |  |  |
| MIRT006689 | *hsa-miR-17-5p* | *RND3* |  |  | ✓ | ✓ |  |  |  |  |
| MIRT006690 | *hsa-miR-103a-3p* | *DAPK1* | ✓ | ✓ |  | ✓ |  | ✓ |  |  |
| MIRT006691 | *hsa-miR-107* | *DAPK1* | ✓ | ✓ |  |  |  | ✓ | ✓ |  |
| MIRT006692 | *hsa-miR-107* | *KLF4* | ✓ | ✓ | ✓ | ✓ | ✓ | ✓ | ✓ | ✓ |
| MIRT006698 | *hsa-miR-499a-5p* | *FOXO4* | ✓ |  | ✓ | ✓ | ✓ |  |  |  |
| MIRT006699 | *hsa-miR-499a-5p* | *PDCD4* | ✓ | ✓ |  | ✓ |  | ✓ | ✓ |  |
| MIRT006785* | *hsa-miR-497-5p* | *IGF1R* | ✓ | ✓ |  | ✓ | ✓ | ✓ | ✓ | ✓ |
| MIRT006785* | *hsa-miR-497-5p* | *IGF1R* |  | ✓ | ✓ | ✓ | ✓ |  |  |  |
| MIRT006825* | *hsa-miR-106a-5p* | *TGFBR2* | ✓ | ✓ | ✓ | ✓ | ✓ | ✓ | ✓ | ✓ |
| MIRT006825* | *hsa-miR-106a-5p* | *TGFBR2* | ✓ |  |  |  |  | ✓ | ✓ | ✓ |
| MIRT006825* | *hsa-miR-106a-5p* | *TGFBR2* |  |  |  |  |  | ✓ | ✓ |  |
| MIRT006825* | *hsa-miR-106a-5p* | *TGFBR2* | ✓ | ✓ |  |  |  | ✓ | ✓ | ✓ |
| MIRT006942* | *hsa-miR-139-5p* | *IGF1R* | ✓ |  | ✓ | ✓ | ✓ | ✓ | ✓ | ✓ |
| MIRT006942* | *hsa-miR-139-5p* | *IGF1R* |  |  |  |  |  |  |  |  |
| MIRT006975 | *hsa-miR-148a-3p* | *BCL2* |  | ✓ | ✓ | ✓ | ✓ | ✓ | ✓ | ✓ |
| MIRT007184 | *hsa-miR-429* | *SOX2* | ✓ | ✓ |  | ✓ | ✓ | ✓ | ✓ | ✓ |
| MIRT007349 | *hsa-miR-362-3p* | *E2F1* | ✓ | ✓ | ✓ |  | ✓ | ✓ |  | ✓ |
| MIRT007350 | *hsa-miR-362-3p* | *USF2* | ✓ | ✓ | ✓ | ✓ | ✓ | ✓ | ✓ | ✓ |
| MIRT007351 | *hsa-miR-362-3p* | *PTPN1* | ✓ |  |  | ✓ |  | ✓ | ✓ | ✓ |
| Here, the asterisk (*) denotes that under the same miRTarBase ID, a single miRNA can interact with multiple regions on the same target mRNA. | | | | | | | | | | |

Recall Table:

| Model_name | miRTarBase datasets (201 samples) |
| --- | --- |
| CIRNN | 0.647 |
| PmliPred | 0.647 |
| PmliHFM | 0.577 |
| LncMirNet | 0.701 |
| PreMLI | 0.716 |
| Gra-CRC-miRTar (GCN) | 0.746 |
| Gra-CRC-miRTar (GAT) | 0.726 |
| Gra-CRC-miRTar (GIN) | **0.786** |
