## Supplementary for "Gra-CRC-miRTar: The pre-trained nucleotide-to-graph neural networks to identify potential miRNA targets in colorectal cancer": Supplement 3.docx

| Model Name | Encoding method | Classifier | Details |
| --- | --- | --- | --- |
| CIRNN | Combining one-hot encoding, The k-mer frequency, GC content, number of base pairs and minimum free energy (MFE). | CNN-IndRNN | CIRNN is a deep learning model developed to predict interactions between plant miRNA and lncRNA. It employs CNN to investigate the functional attributes of gene sequences and uses Independently Recurrent Neural Networks (IndRNN) to capture the sequence features' representation. |
| PmliPred | Combining one-hot encoding, The k-mer frequency, GC content, number of base pairs and minimum free energy (MFE). | CNN-BRU | PmliPred is a deep learning model for plant miRNA-lncRNA interaction prediction, utilizing a CNN-BiGRU architecture. It encodes RNA sequences through one-hot encoding alongside manually extracted features, incorporating fuzzy decision-making to enhance prediction accuracy. |
| PmliHFM | High-order one-hot encoding. | CNN + stacked encoder-decoder | PmliHFM is a deep learning model designed for the prediction of plant miRNA-lncRNA interactions. It utilizes a hybrid feature mining network architecture that incorporates both a CNN and a stacked encoder-decoder. It employs different encoding methods for miRNA sequence and lncRNA sequence due to their different sequence lengths. |
| LncmirNet | Including four sequence-based features in the encoding: K-mer features, CTD features, Doc2vec features, Graph embedding generated by Role2vec. | CNN | LncMirNet is a deep learning model designed for predicting miRNA-lncRNA interactions, leveraging a CNN architecture. It innovatively processes RNA sequences by encoding them with four types of sequence-based features, including graph embedding features. The model employs a histogram-dd method to integrate these features into a matrix for the CNN's learning pattern. |
| preMLI | Rna2vec | CNN-BRU (Attention mechanism includes) | PreMLI is a deep learning model designed for predicting miRNA-lncRNA interactions, utilizing a CNN-BiGRU architecture. It obtains low-dimensional feature vectors from lncRNA and miRNA sequences through the rna2vec method. Additionally, an attention mechanism is integrated to focus on key features within these vectors. |
